# PARP12-catalyzed mono-ADP-ribosylation of Golgin-97 controls the transport of E-cadherin

**DOI:** 10.1101/2020.05.05.078097

**Authors:** Giovanna Grimaldi, Laura Schembri, Matteo Lo Monte, Daniela Spano, Rosaria Di Martino, Andrea R Beccari, Carmen Valente, Daniela Corda

## Abstract

ADP-ribosylation is a post-translational modification involved in physiological and pathological events catalyzed by Poly-ADP-Ribosyl-Polymerase (PARP) enzymes. Substrates of this reaction have been identified by mass-spectrometry, but the definition of PARPs-regulated cellular functions remains scarce. Here, we have analyzed the control of intracellular membrane traffic by the mono-ADP-ribosyl-transferase PARP12, motivated by its localization at the *trans*-Golgi network. By using bioinformatics, mutagenesis and cell biology approaches we identified Golgin-97, a protein regulating exocytosis, as a PARP12-specific substrate. Mono-ADP-ribosylation of Golgin-97 residues E558-E559-E565 is required for supporting traffic from the *trans*-Golgi network to the plasma membrane. This step is halted when PARP12 is deleted or when the Golgin-97 ADP-ribosylation-defective mutant is expressed. Under these conditions E-cadherin, whose transport is controlled by Golgin-97, does not reach the plasma membrane but accumulates in a *trans*-Golgi proximal compartment. Thus, we demonstrate that the ADP-ribosylation of Golgin-97 is required for E-cadherin exocytosis and thus this event may regulate the sorting of exocytic carriers as well as epithelial-to-mesenchymal transition.

## Introduction

The poly-ADP-ribose polymerase (PARP) family includes 17 members, that share a conserved catalytic domain functional to the transfer of ADP-ribose (ADPr) from NAD^+^ to specific amino-acid residues in target proteins; the addition of a single ADPr is known as mono-ADP-ribosylation (MARylation), while the addition of multiple ADPr units (by elongation or branching) is known as poly-ADP-ribosylation (PARylation) (Otto et al., 2005). The majority of PARPs have MARylation activity, while only PARP1, −2 and −5 act as polymerases (Vyas et al., 2014). The PARPs’ catalytic pocket is characterized by a conserved Histidine-Tyrosine-Glutamate (HYE) triad, also present in the ADP-ribosyl-transferase (ART) fold of diphtheria toxin; thus, PARPs are also named ADP-ribosyltransferases diphtheria toxin-like (ARTD) (Hottiger et al., 2010). PARPs are localized at diverse subcellular compartments (*e*.*g*., nucleus, endoplasmic reticulum, Golgi complex *etc*.), hinting at potential, diverse biological functions they contribute to (Cohen and Chang, 2018; Corda and Di Girolamo, 2003; Di Girolamo et al., 2005; Dolle et al., 2010; Gupte et al., 2017; Hassa and Hottiger, 2008).

MARylation was identified decades ago as a protein, post-translational modification (PTM) exploited by bacterial toxins to promote bacterial pathogenesis in host cells (Gill et al., 1969; Moss and Vaughan, 1977). While the occurrence of this protein PTM in mammalian cells has soon become evident (Dani et al., 2011; Jones and Baird, 1997; Lupi et al., 2000; Lupi et al., 2002; Moss et al., 1999; Okazaki and Moss, 1996; Seman et al., 2004) only recently, with the definition of the different PARPs (and of their intrinsic enzymatic activities) the physiological functions the MARylating PARPs may regulate are emerging (Cohen and Chang, 2018; Gupte et al., 2017). Indeed, MARylation as the other PTMs [over 200 so far identified; (Mann and Jensen, 2003)] are expected to convey to proteins the ability to signal in diverse regulatory cascades, favoring diversified, and rapid intracellular communication.

So far, ADP-ribosylation has been mainly related to stress conditions, as exemplified by the role of PARP1-mediated PARylation during DNA-damage response (Luo and Kraus, 2012), or PARP5, −12 and −13 role in stress granule formation (Catara et al., 2017; Leung, 2014; Leung et al., 2011), or PARP16 in the unfolded protein response (Di Paola et al., 2012; Jwa and Chang, 2012). In oxidative stress, PARP-derived Poly-ADP-ribose (PAR) forms the backbone essential in the formation of stress granules (SGs), non-membranous structures assembled following different kind of stresses (Buchan and Parker, 2009; Kedersha and Anderson, 2002). Importantly, PAR has also a signaling role during stress response; we have recently reported that PARP1-produced PAR is the signal leading to PARP12 translocation from the Golgi complex (its localization under steady-state conditions) to SGs (Catara et al., 2017). This translates in the block of anterograde transport, from the *trans*-Golgi network (TGN) to the plasma membrane (PM), determining the stall of intracellular membrane transport as part of the stress response and underlining a key role of PARP12 in controlling exocytic pathway (Catara et al., 2017; Grimaldi et al., 2019; Grimaldi and Corda, 2019).

The involvement of ADP-ribosylation in intracellular membrane transport is indeed emerging as a novel function mediated by the PARPs, with particular reference to PARP5 and −12 (Grimaldi and Corda, 2019). PARP5 (named also tankyrase) - the other Golgi complex-localized PARP - is known to regulate the delivery of the glucose transporter GLUT4 from the TGN to glucose-storage vesicles and thus to the PM (Chi and Lodish, 2000; Guo et al., 2012; Su et al., 2018; Yeh et al., 2007). Differently, PARP12 has mainly been related to viral infections add new refs (Atasheva et al., 2012; Atasheva et al., 2014; Li et al., 2018; Welsby et al., 2014) or as a component of SGs (Catara et al., 2017; Leung et al., 2011). The reported PARP12 participation in intracellular membrane transport points at a novel function for PARP12 that deserves attention.

Here we identify Golgin-97 as the substrate of PARP12-catalyzed MARylation and define the residues targets of this modification; this is so far the only known PTM of Golgin-97 controlling its activity. Golgins are proteins that localize at different stacks of the Golgi complex, typically attached to the membrane at their carboxyl termini *via* a trans-membrane domain or a domain that binds a small GTPase of the Rab or Arf families (Cheung and Pfeffer, 2016; Witkos and Lowe, 2015); here, Golgins contribute to the maintenance of the organelle structure and to the regulation of vesicular transport, mostly as vesicle tethers (Lowe, 2019; Muschalik and Munro, 2018). In addition, Golgin-97 and Golgin-245 are also involved in the exocytosis of E-cadherin and TNFα, respectively (Lieu et al., 2008; Lock et al., 2005). However, the molecular mechanisms underlying these Golgins’ functions remain to be defined (Lieu et al., 2008; Lock et al., 2005).

In this study, we demonstrate that PARP12-mediated MARylation of Golgin-97 regulates the transport of E-cadherin; the absence of the enzyme, as well as the lack of ADP-ribosylated Golgin-97, impinges on the correct arrival of E-cadherin to the PM. Thus, ADP-ribosylation of Golgin-97 is a required PTM to control Golgin-97-driven exocytosis. This event, as discussed, can play a role in determining transport-carrier sorting to the PM and, by affecting E-cadherin transport, be part also of the EMT control.

## Results

### The exocytosis of E-cadherin is specifically controlled by PARP12 and Golgin-97

We have reported that PARP12, that is localized at the TGN, guarantees the correct export of the basolateral cargo vesicular stomatitis virus G protein (VSVG) from the TGN to the PM (Catara et al., 2017). The exit of basolateral cargoes from the TGN occurs in different carriers, possibly exploiting different sorting mechanisms (Boncompain et al., 2012). For instance, the exit of the VSVG cargo from the TGN occurs in tubulovesicular carriers (De Matteis and Luini, 2008; Pagliuso et al., 2016; Polishchuk et al., 2003; Valente et al., 2012), similar to those used by the cargo E-cadherin (Boncompain et al., 2012), while other cargoes, exemplified by the TNFα, use vesicular carriers to exit this compartment (Boncompain et al., 2012). To better understand the role of PARP12 in regulating cargo exiting from the TGN, we analyzed its role in the export of two different basolateral cargoes, E-cadherin and TNFα, as examples of the two different sorting routes proposed so far (Boncompain et al., 2012).

To this aim, we took advantage of the well-established synchronizable Retention Using Selective Hook (RUSH) transport system (Boncompain et al., 2012) (see methods). In this system, a reporter protein (*i*.*e*. the cargo) is fused with streptavidin-binding peptide (SBP); its interaction with a hook protein fused with streptavidin is responsible for reporter retention at the donor compartment. Biotin addition competitively displaces this interaction, thus enabling the reporter to be synchronously released from the donor to its acceptor compartment (PM in our case). HeLa cells, devoid or not of PARP12 expression, were transiently transfected with SBP-EGFP-E-cadherin (Fig 1A) or TNFα-SBP-EGFP (Fig 1B) and exiting of cargoes from the TGN towards the PM evaluated, by quantifying the amount of cargo at the Golgi complex after 60 min of biotin addition (Fig 1C, D). The transient depletion of PARP12 in HeLa cells alters the exit from the TGN, as indicated by the increased cargo entrapped on the Golgi complex (Fig 1A and C, see methods), while not affecting the exocytosis of TNFα (Fig 1B and D).

**Fig 1.**
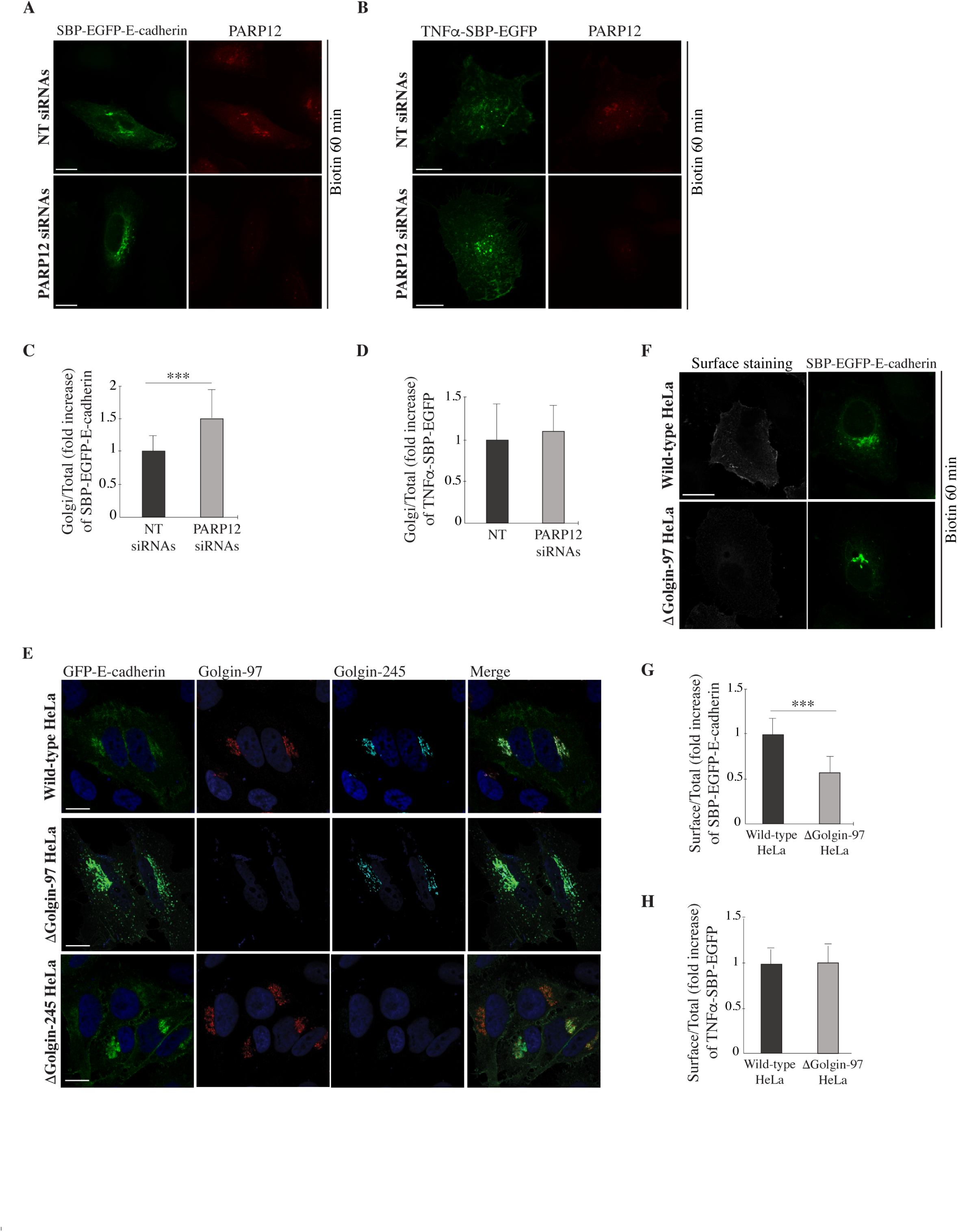
E-cadherin exocytosis is controlled by PARP12 and Golgin-97. HeLa cells expressing the indicated RUSH-based (SBP) constructs, showing increased amount of EGFP-E-cadherin (green) (**A**), but not EGFP-TNFα (green) (**B**), at the Golgi complex after 60 min of biotin addition, in the absence of PARP12 (PARP12 siRNAs). Quantifications are shown in graphs (**C-D**). (**E)** Representative confocal images of GFP-E-cadherin (green) localization in wild-type, ΔGolgin-97 and ΔGolgin-245 cells, showing E-cadherin at the PM in control and ΔGolgin-245, but not in ΔGolgin-97, cells. Golgin-97 is shown in red; Golgin-245 is in cyan. (**F**) Representative confocal images of the E-cadherin pool at the PM (surface staining, gray) of synchronized RUSH-based E-cadherin (green) in wild-type and ΔGolgin-97 cells; the quantifications show a reduction of the E-cadherin (**G**), but not of TNFα (**H**), staining at the PM in the absence of Golgin-97. n = 30 cells from three different experiments ± SD. ***p < 0.001; calculated by Student’s t test. Scale bars, 10 µm.

E-cadherin and TNFα transport is specifically controlled by two distinct Golgins, Golgin-97 and Golgin-245, respectively (Lieu et al., 2008; Lock et al., 2005). We assessed the specificity of Golgin-97 in E-cadherin exocytosis by analyzing the localization of overexpressed GFP-tagged E-cadherin in HeLa cells deleted of Golgin-97 [by CRISPR-Cas9 technology, referred as ΔGolgin-97 cells, (Shin et al., 2017; Wong and Munro, 2014)] or, as a negative control, deleted of Golgin-245 [ΔGolgin-245 cells, (Shin et al., 2017)], and compared them to their wild-type HeLa cells. While E-cadherin clearly stained the PM (thus forming adherent junctions) in both wild-type and ΔGolgin-245 cells, the absence of Golgin-97 caused a defect in E-cadherin localization, with the cargo being trapped in intracellular E-cadherin–containing transport carriers (Fig 1E) localized in a post-TGN/Rab11-positive intermediate compartment (Fig EV1). By analyzing HeLa cells lacking Golgin-245 we could exclude a redundant role of Golgin-245 in E-cadherin exocytosis (Fig 1E), at variance with the similar function as tethers in endosome-to-Golgi trafficking exerted by both Golgin-97 and −245 (Shin et al., 2017; Wong et al., 2017; Wong and Munro, 2014).

The specificity of Golgin-97 in controlling E-cadherin exocytosis was further analyzed in wild-type and ΔGolgin-97 HeLa cells using the RUSH system, by evaluating the E-cadherin pool at the PM (see methods). Absence of Golgin-97 specifically reduced the arrival of E-cadherin at the PM (Fig 1F and G), while not affecting the transport of TNFα (Fig 1H), similarly to the data observed upon PARP12 depletion (Fig 1A-D).

As previously reported by us (Catara et al., 2017), lack of PARP12 prevents VSVG exocytosis, thus the potential contribution of Golgin-97 in this transport step using ΔGolgin-97 HeLa cells was analyzed by means of the VSVG-based transport assay (Catara et al., 2017; Valente et al., 2012). Briefly, cells were infected with VSV and incubated at 40 °C to first accumulate the protein in the endoplasmic reticulum. Cells were then shifted to 20 °C, a temperature that allows the exit of the cargo proteins from the endoplasmic reticulum and the arrival to, but not the exit from, the TGN. The temperature was finally shifted to 32 °C, and transport from the TGN to the plasma membrane was monitored by immunofluorescence (Fig EV2). Compared to wild-type HeLa cells, absence of Golgin-97 increases the percentage of cells showing VSVG at the Golgi complex after 30 min of the temperature block release (Fig EV2), demonstrating a role of Golgin-97 in VSVG exocytosis. Collectively, these data indicate that PARP12 and Golgin-97 cooperate in the regulation of the exocytosis of the same class of basolateral cargoes, thus possibly contributing to determine sorting.

### Golgin-97 is a PARP12 substrate

The PARP12-mediated modulation of E-cadherin exocytosis (Fig 1) could require PARP12-catalyzed modification of Golgin-97. To investigate this aspect, GFP-tagged E-cadherin localization was evaluated in wild-type HeLa cells treated with the general PARP inhibitor PJ34 (50 µM; see methods). This, at variance with untreated cells, caused a mis-localization of E-cadherin that was accumulated in spots reminiscent of transport carriers (Fig 2A). With synchronized E-cadherin (using the RUSH system; see methods) PJ34 treatment delayed its exiting from the TGN (Fig 2B, C), thus suggesting an involvement of PARP12 enzymatic activity in E-cadherin transport, possibly through Golgin-97 modification. Thus, Golgin-97 was transiently overexpressed in HeLa cells and the total cell lysates analyzed using the Af1521 *macro* domain based pull-down assay (Dani et al., 2009; Grimaldi et al., 2018), which detected ADP-ribosylated Golgin-97 (Fig 2D), but not Golgin-245 (used as control of specificity, Fig 2E); this indicates that only Golgin-97 was modified under these experimental conditions (Fig 2D; see methods). Golgin-97 MARylation was also analyzed after PARP12 depletion (by specific siRNAs: about 50% enzyme reduction) and resulted reduced by about 50% (Fig 2F), indicating that Golgin-97 is a substrate of PARP12 activity. The potential contribution of PARP5 [also localized at the Golgi complex (Chi and Lodish, 2000)] in this PTM was evaluated in cells treated for 2 h with PJ34 (50 µM) or with the PARP5-specific inhibitor IWR1 (25 µM): Golgin-97 was not ADP-ribosylated in the presence of PJ34 while it was in the presence of IWR1, pointing at the specificity of the PARP12-dependent modification (Fig 2G).

**Fig 2.**
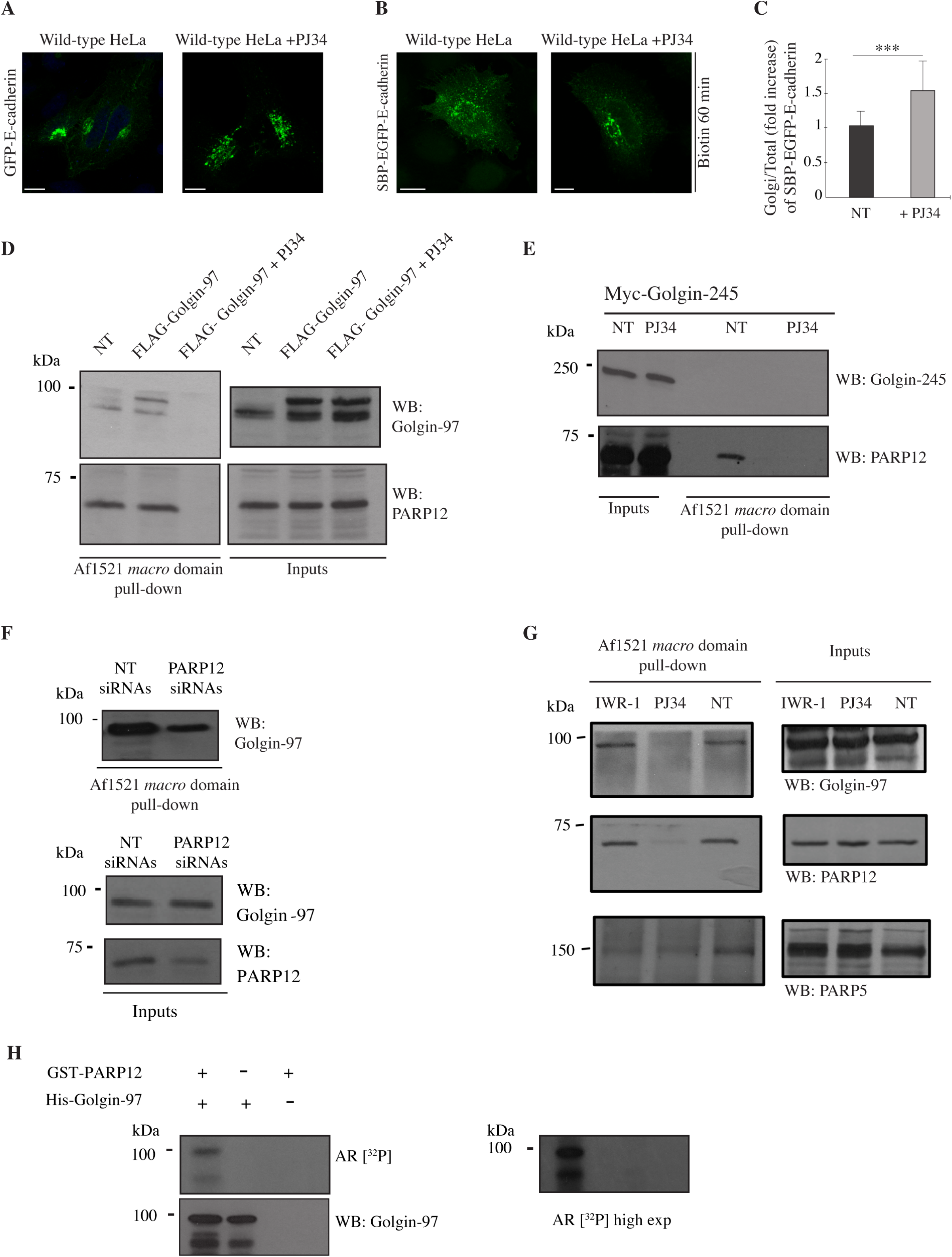
Golgin-97 is a PARP12 MARylation substrate. Representative confocal images of GFP-E-cadherin (green) (**A**) or RUSH-based (SBP) EGFP-E-cadherin (**B**) localization in wild-type HeLa cells treated or not with PJ34 (50 µM, 2 h). (**C**) Quantification of E-cadherin at the Golgi complex, showing an increase in cells treated with PJ34. (**D, E**) Af1521 *macro* domain based pull-down assay of total cell lysates obtained from cells untreated (NT) or treated with the general PARP inhibitor PJ34 (50 µM, 2 h) showing detection of MARylated, FLAG-tagged Golgin-97 (D), but not of Myc-tagged Golgin-245 (E). (**F**) HeLa cells depleted of PARP12 (PARP12 siRNAs) or not (NT siRNAs), and (**G**) untreated (NT) or treated with PJ34 (50 µM, 2 h) or with IWR1 (25 µM, 2 h) showing the PARP12-dependent MARylation of Golgin-97. PARP12 and PARP5 were detected as internal controls (G). (**H**) *In vitro* MARylation assay using GST-tagged purified PARP12 catalytic fragment and His-tagged purified Golgin-97, in presence of [^32^P]-NAD^+^, detected by autoradiography (AR [^32^P]). Lower panel shows total levels of Golgin-97. n = 30 cells from three different experiments ± SD. ***p < 0.001; calculated by Student’s t test. Scale bars, 10 µm.

Golgin-97 MARylation was then analyzed in *in vitro* assays, using purified full-length Golgin-97 and PARP12-catalytic domain, in the presence of ^32^P-NAD^+^ (see methods). Golgin-97 was modified under these conditions, indicating that it is a specific, direct substrate of PARP12-dependent ADP-ribosylation (Fig 2H).

Altogether, these data demonstrate that Golgin-97 is specifically MARylated by PARP12 both *in vitro* and in intact cells, thus suggesting that the effects of the impaired exocytosis reported above may be related to this specific PTM.

### Identification of the Golgin-97-modified residue(s)

Having defined Golgin-97 as a PARP12 specific substrate, we investigated the specific residue(s) target(s) of this PTM. To this purpose and based on our previous data demonstrating that PARP12 modifies acid residues (Catara et al., 2017), we exploited *ADPredict*, an in-house developed bioinformatic tool (freely accessible online at *www.adpredict.net)* for the prediction of the acidic residues (aspartic and glutamic acids, D and E) most prone to be ADP-ribosylated (Lo Monte et al., 2018). The predictions highlighted two main regions enriched in putative sites of ADP-ribosylation: a first one, spanning from position 367 to 405, and a second one from 548 to 585 (see Appendix Table S1, Fig EV3). On these premises, we set-up a multiple mutant strategy, consisting of three separated multiple point mutants, from here on addressed as Mut A: E381Q-E386Q-E393Q; B: E558Q-E559Q-E565Q; C: E579Q-E585Q (see Appendix Figure S1). Conservative mutations were introduced (exchanging acidic residues with their relative amide, E to Q) and mutants cellular localization analyzed by immunofluorescence (see methods). All the mutants localized at the TNG, similarly to the wild-type protein (Fig 3A). This was more evident after nocodazole-induced disruption (33 μM for 3 h) of the Golgi ribbon into smaller structures, the “mini-stacks” (Fig 3B), suitable to precisely determine protein localization across the Golgi stacks (Beznoussenko et al., 2014).

**Fig 3.**
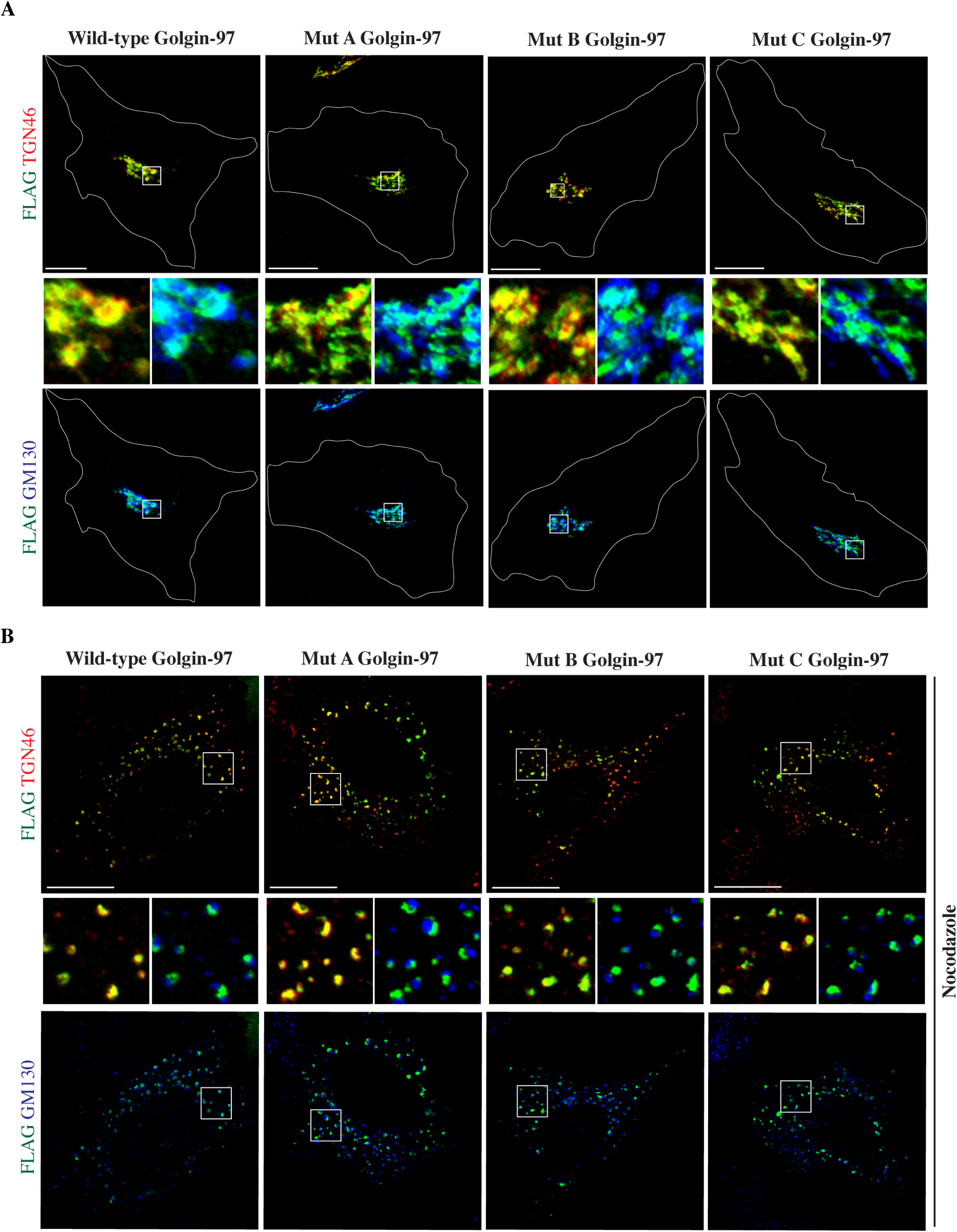
Subcellular localization of Golgin-97 multiple point mutants. HeLa cells were transfected as indicated, and treated with (**A**) vehicle alone or (**B**) nocodazole (33 μM, 3 h), showing a correct TGN-localization of FLAG-tagged (green) Golgin-97 point mutants as for the wild-type Golgin-97 with the TGN marker TGN46, but not with the cis-Golgi marker GM130 (blue). Insets: enlarged view of merged signals at the Golgi membranes. Scale bars, 10 μm. See also Fig EV3 and Appendix Table S1 for the identification of the residues by the ADPredict.

The ADP-ribosylation of the Golgin-97 point mutants was thus analyzed in HeLa cells, using the Af1521 *macro* domain-based pull-down assay on total cell lysates overexpressing the different mutants or the wild-type protein (see methods). Of note, while the wild-type Golgin-97 was recognized by the Af1521 *macro* domain, the binding of mutants A, B and C was strongly reduced (Fig 4A). Similar results were obtained using ΔGolgin-97 HeLa cells (Fig 4B). To further demonstrate these residues as targets of MARylation, *in vitro* ADP-ribosylation assays were performed using purified full-length Golgin-97 (wild-type and the different mutants) and the PARP12-catalytic domain. The pool of modified Golgin-97 was recovered by using Af1521 *macro* domain as a bait (see methods) and the percentage of MARylated protein was calculated as ratio between Af1521-bound Golgin-97 and total Golgin-97 (*i.e*. input material, see Fig 4C). The results obtained confirm the data deriving from cell lysates (Fig 4A, B), demonstrating that the three multiple point mutants generated are ADP-ribosylation defective mutants (Fig 3, 4). Golgin-97 harbors two main domains (see Appendix Figure S1): the GRIP domain - responsible for its targeting to the Golgi complex - and the coiled-coil domain, involved in protein-protein interactions [e.g. with FIP1/RCP or Rabs (Jing et al., 2010; Wong et al., 2017)]. The residues identified as targets of MARylation are all included in the coiled-coil domain of Golgin-97, suggesting a role in the regulation of protein interactions.

**Fig 4.**
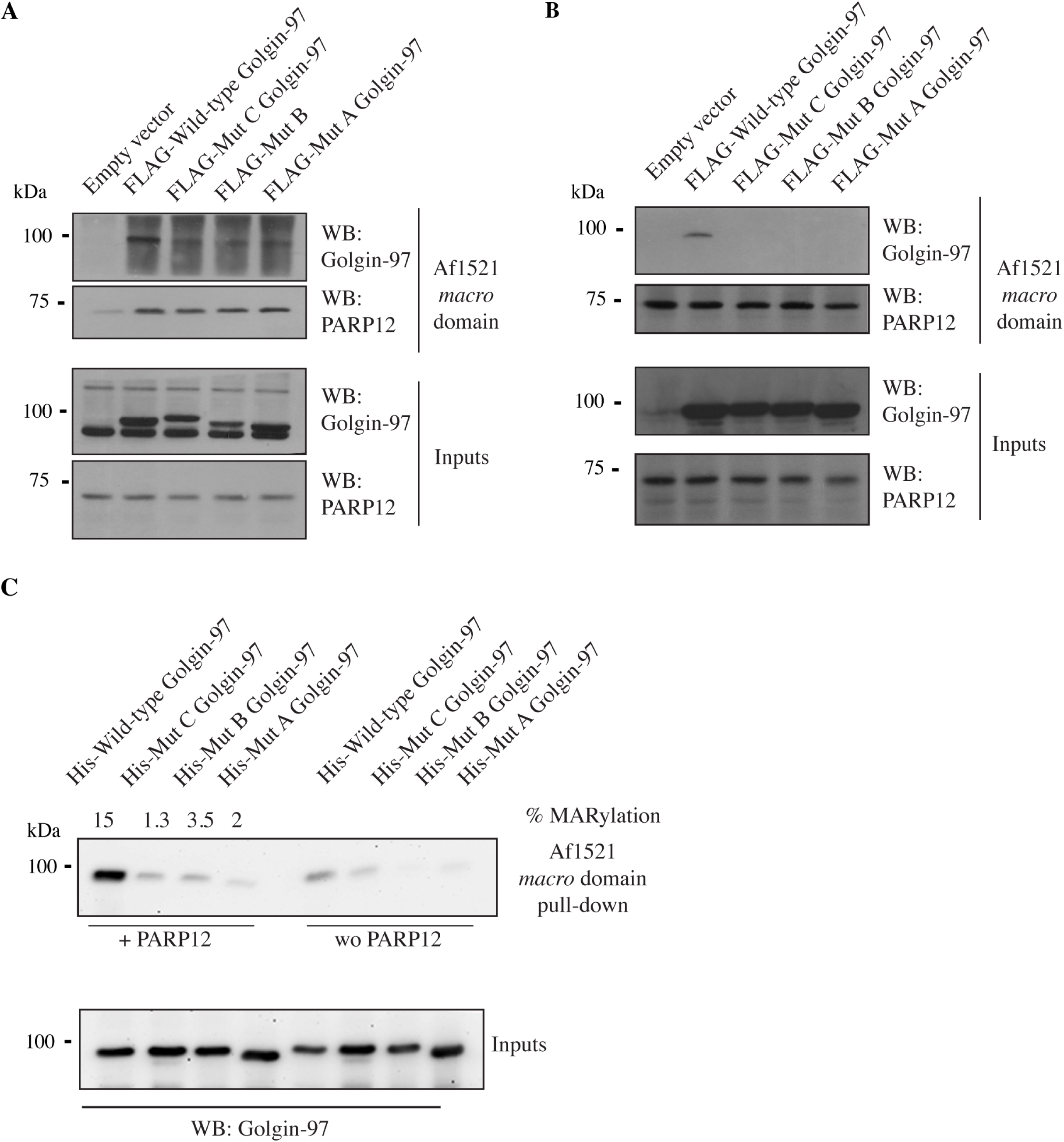
Validation of Golgin-97 MARylation-defective mutants. (**A, B**) Af1521 *macro* domain pull-down assay of lysates obtained from HeLa cells transiently transfected with the full-length Golgin-97 or with its multiple point mutants. The bound proteins were eluted and detected by Western blotting (WB) with anti-Golgin-97 antibody. MARylation of PARP12 detected as internal control. (**C**) *In vitro* MARylation assay using GST-tagged purified PARP12 catalytic fragment and His-tagged purified Golgin-97 (wild-type and multiple point mutants) was followed by the Af1521 *macro* domain pull-down assay to recover the pool of MARylated Golgin-97. The bound proteins were eluted and detected by Western blotting (WB) with anti-Golgin-97 antibody. Signals deriving from samples incubated with the reaction buffer alone (wo PARP12) were considered as background signals. Lower panel shows input materials. Molecular weight standards (kDa) are indicated on the left of the panel.

Based on these data we concluded that the coiled-coil domain of Golgin-97 includes the residues targets of the ADP-ribosylation catalyzed by PARP12.

### ADP-ribosylated Golgin-97 is required for a correct export of E-cadherin

To assess the significance of MARylation in the export of E-cadherin, GFP-tagged E-cadherin localization was evaluated in wild-type or ΔGolgin-97 HeLa cells, the latter further transiently transfected with the wild-type Golgin-97 or with its ADP-ribosylation defective mutants (see methods). Ectopically added wild-type Golgin-97 rescued the localization defect observed in ΔGolgin-97 HeLa cells, resulting in a clear PM localization of the cargo (E-cadherin at PM, forming adherent junction; Fig 5A), similarly to the parental cell line (wild-type HeLa; Fig 5B); differently, mutant B was unable to recover the defective phenotype, and showed the same accumulation of GFP-tagged E-cadherin in perinuclear carriers (Fig 5A). Mutants A and C instead recovered the defect as the wild-type Golgin-97 (Fig 5A). These data indicate that Golgin-97 is ADP-ribosylated on different residues regulating multiple functions; the ADP-ribosylation occurring at E558-E559-E565 (mutated in Mut B) specifically modulates Golgin-97 role in E-cadherin exocytosis.

**Fig 5.**
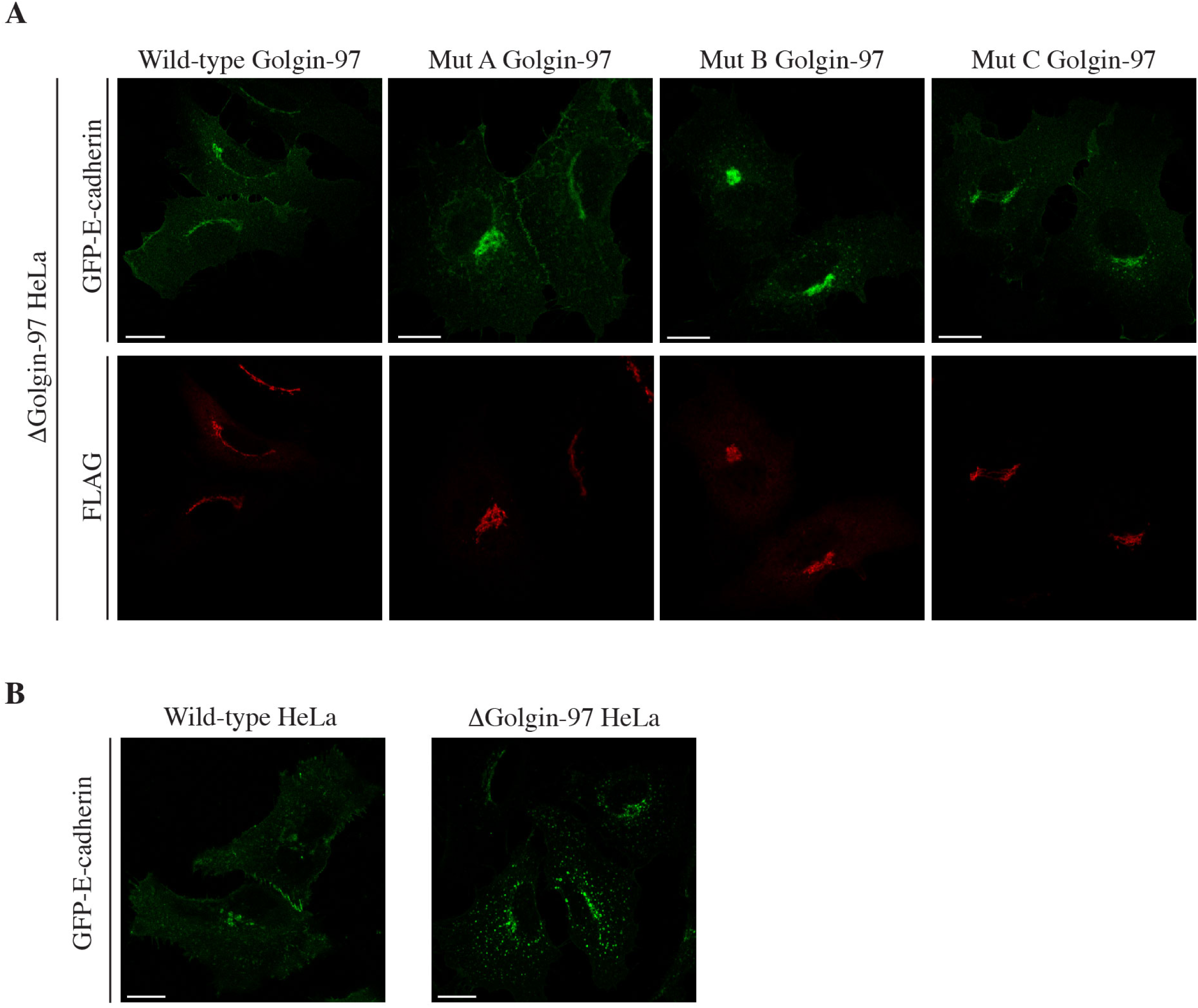
MARylated Golgin-97 is required for a correct E-cadherin localization. Representative confocal images of **(A)** GFP-E-cadherin (green) localization in ΔGolgin-97 cells showing the rescue of E-cadherin localization and formation of adherent junctions with FLAG-tagged wild-type Golgin-97 or Mut A and C, but not with Mut B. **(B)** GFP-E-cadherin localization in wild-type and ΔGolgin-97 HeLa cells is reported for comparison. Scale bars, 10 µm.

These results were further substantiated by monitoring the GFP-tagged E-cadherin transport in live imaging experiments performed in wild-type or ΔGolgin-97 HeLa cells, the latter further transiently transfected with the wild-type Golgin-97 or with its ADP-ribosylation defective mutant B, which showed, respectively, the rescuing or impairment of the E-cadherin transport, as reported above (see methods and Movies EV 1-4, Fig EV4).

Collectively, these results are in line with the requirement of Golgin-97 MARylation in controlling E-cadherin exocytosis.

### PARP12-mediated ADP-ribosylation of endogenous Golgin-97 in MCF7 cells

The relevance of PARP12 activity in E-cadherin exocytosis was substantiated by evaluating the E-cadherin localization in epithelial MCF7 cells (that endogenously express E-cadherin) by immunofluorescence. PARP12-depleted cells did not show E-cadherin staining at the PM, and were not able to form adherent junctions (Fig 6A). A similar phenotype was observed in MCF7 cells transiently depleted of Golgin-97 (Fig EV5).

**Fig 6.**
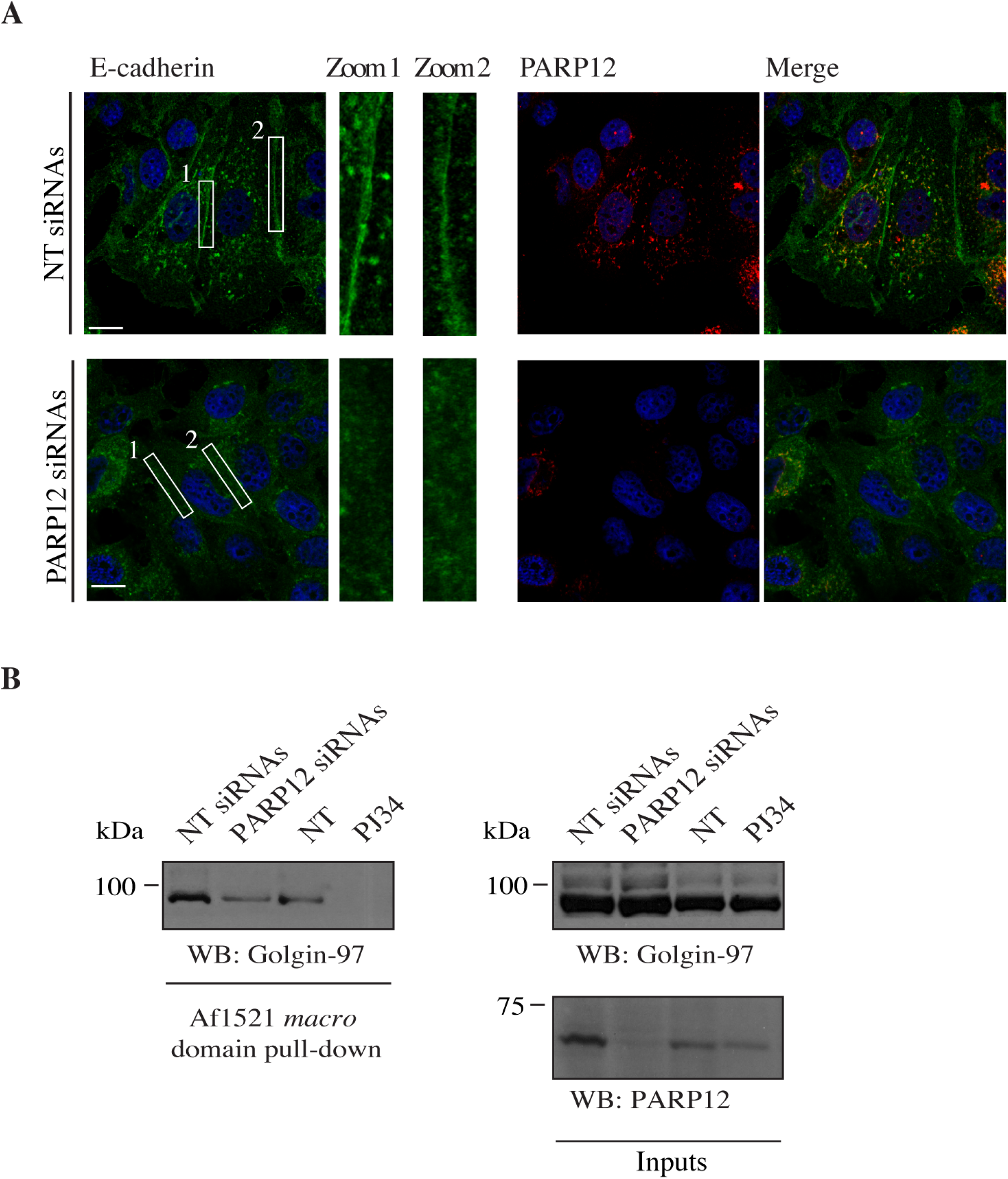
Endogenous E-cadherin localization is controlled by the PARP12 -mediated Golgin-97 MARylation in epithelial MCF7 cells. **(A)** Representative confocal images of the decreased endogenous E-cadherin (green) localization at PM in MCF7 cells transiently depleted of PARP12. Insets: enlarged view of E-cadherin signal at PM, where E-cadherin forms adherent junctions, stabilizing cell-cell adhesions. See also Fig EV5, showing the lack of E-cadherin localization at the PM in MCF7 cells in the absence of Golgin-97. **(B)** Af1521 *macro* domain based pull-down assay of total cell lysates obtained from MCF7 cells depleted of PARP12 (PARP12 siRNAs) or not (NT siRNAs), and from MCF7 untreated (NT) or treated with PJ34 (50 µM, 2 h) showing the PARP12-dependent MARylation of Golgin-97. Molecular weight standards (kDa) are indicated on the left of the panels. Scale bars, 10 µm.

To corroborate our findings, ADP-ribosylation of endogenous Golgin-97 was evaluated using the Af1521 *macro* domain-based pull-down assay on total cell lysates of MCF7 cells at steady state, transiently depleted of PARP12 or treated with the PARP inhibitor PJ34 (50 µM; see methods). Along the same line of evidence deriving from the data on E-cadherin localization reported above (Fig 6A), endogenous Golgin-97 was indeed recognized by the Af1521 *macro* domain; its binding was strongly reduced both in the absence of PARP12 or when its catalytic activity was inhibited (Fig 6B).

Together these data - obtained in a cell system expressing endogenous E-cadherin - reinforce the importance of the PARP12-mediated MARylation of Golgin-97 in controlling E-cadherin localization.

## Discussion

With this study, by demonstrating that the ADP-ribosyltranferase PARP12 catalyzes the MARylation of Golgin-97, and that this PTM is required to guarantee the correct transport of E-cadherin from the TGN to the PM, we identify MARylation as an essential player in the regulation of intracellular membrane transport.

So far, the functions of PARP enzymes were mainly connected to the stress response, while limited examples of their roles in the regulation of physiological processes were available. Among these, of note is the function of PARP5 in mitotic progression (Chang et al., 2005a, 2009; Chang et al., 2005b; Slade, 2019) and in GLUT4 transport (Chi and Lodish, 2000; Guo et al., 2012; Su et al., 2018; Yeh et al., 2007).

With our data we point at the importance of MARylation in controlling the exocytic pathway, as it is regulated by the modified Golgin-97; in addition, they also pinpoint the first PTM that regulates Golgin-97 functions.

Golgins are proteins associated to the Golgi complex, with roles in intracellular membrane transport and maintenance of the Golgi complex structure (Kulkarni-Gosavi et al., 2019; Muschalik and Munro, 2018). The molecular mechanisms underlying these functions are just starting to emerge. For example, the tethering of specific vesicles by Golgin-97 and −245 is determined by their interaction with TBC1D23 (a catalytically inactive Rab GAP) and FAM21, a subunit of the WASH complex (Shin et al., 2019; Shin et al., 2017; Wong et al., 2017; Wong and Munro, 2014). The tethering function of Golgins may be inhibited by the phosphorylation of their interactors as in the case of the CDK1-mediated phosphorylation of GM130 during mitosis, that prevents its binding to p115-containing vesicles (Lowe et al., 1998; Nakamura et al., 1997). In the case of Golgin-97 no evidence has been reported to our knowledge of a regulatory PTM other than the present report. Other PTM may however be expected, but it stays that the MARylation of Golgin-97 is an essential PTM for the exocytosis of E-cadherin.

Our findings include the definition of the residues target of MARylation (Fig 3, 4), obtained by using the bioinformatic tool we recently developed (Lo Monte et al., 2018) that, now successfully applied also to Golgin-97, represents an important asset to study D and E MARylation.

Indeed, we can conclude that the MARylation of E558-E559-E565 sites determines the Golgin-97-dependent, correct delivery of E-cadherin to the PM. Of interest, not all the ADP-ribosylation residues identified supported E-cadherin transport (Fig 5), suggesting that they are involved in different Golgin-97 functions (possibly through the correct interaction with diverse complementary proteins).

The identification of the residues E558-E559-E565 is in line with our data indicating that PARP12 MARylation occurs mainly on acidic residues [as it is sensitive to hydroxylamine treatment; (Catara et al., 2017)]. It is worth mentioning that serines (S) are now well documented residues modified by PARP1 under stress conditions (Abplanalp et al., 2017; Bonfiglio et al., 2017; Fontana et al., 2017; Larsen et al., 2018; Palazzo et al., 2018). While our data support E residues as the main Golgin-97-residues modified by PARP12, we cannot exclude that S may also be modified by other PARPs or under different experimental conditions. A possibility is that different modified residues determine the interaction with the complex of proteins needed for the specific signaling pathways or cell process (*e.g*., stress conditions *vs* physiological conditions).

A novel aspect emerging from our data is the cargo specificity provided by the PARP12-mediated ADP-ribosylation of Golgin-97 in regulating exocytosis. This aspect gains particular interest when considering the molecular mechanisms dictating different sorting pathways. E-cadherin (as well as VSVG) and TNFα are known to exit the TGN with different kinetics and in morphologically distinct carriers, *i.e*. tubular or granule-like post-Golgi carriers, respectively (Boncompain et al., 2012). These phenotypes have been proposed to be supported by a sequential segregation of the two cargoes both intra- and post-Golgi (Boncompain et al., 2012), but how this segregation occurs at the molecular level is still not clear.

The sequential segregation hypothesis has been also proposed in a different study, that analyzes the export of endolysosomal proteins out of the TGN (Chen et al., 2017). The authors described two post-Golgi carrier populations, tubular carriers for transferrin receptor and LAMP1 (which also contain VSVG when co-transported) and vesicular carriers for cationic-dependent mannose-6-phosphate receptor (CD-MPR).

Based on these studies, one can assume that cargoes exiting the TGN in tubular carriers depend on Golgin-97 (as in the specific case of E-cadherin and VSVG), while those cargoes exiting in vesicle-like structures depend on other mechanisms (possibly connected to different Golgins/adaptors).

Considering *i)* the role of PARP12 and Golgin-97 in specifically regulating E-cadherin export (while not affecting TNFα) and *ii)* the two morphologically-different type of carriers containing E-cadherin and TNFα while exiting the Golgi complex (Boncompain et al., 2012), we propose that the PARP12-mediated MARylation of Golgin-97 regulates this sequential sorting, and specifically intervenes on those cargoes exiting the TGN through tubular carriers.

In conclusion, the data reported indicate that PARP12-catalyzed MARylation modulates the function of Golgin-97 (and more in general of a given substrate) and that this PTM accounts for the multiple functions the protein might have. In particular, we have focused on the MARylated Golgin-97-dependent transport of E-cadherin to the PM. Here, E-cadherin mediates the formation of adherent junctions, required to maintain an intact architecture of the normal epithelium (Bruser and Bogdan, 2017). Tumoral cells are characterized by loss of adherent junctions and thus cell polarity, while gaining migratory and invasive properties, an event known as epithelial-to-mesenchymal transition [EMT, (Prieto-Garcia et al., 2017)]. Cadherins dysregulation is part of the EMT (Hanahan and Weinberg, 2011): loss of E-cadherin from the PM is an early indicator of EMT and a marker of poor prognosis in many cancers (Bremnes et al., 2002; Chen et al., 2003; Fan et al., 2019; Sommariva and Gagliano, 2020), making the trafficking of E-cadherin an area of great interest for the discovery of new mechanisms that modulate cell adhesion (Cadwell et al., 2016).

An altered trafficking of E-cadherin impinges on the stability of the adherent junctions, as already reported for defective endosomal trafficking of E-cadherin (Palacios et al., 2005). Here we report on Golgin-97 MARylation as a new regulator of E-cadherin exocytosis, an event with consequences on adherent junction stability. We envision that a hampered Golgin-97 MARylation associated to defective E-cadherin transport contributes to the loss of adherent junctions (see Figs 5, 6) and thus EMT induction. Considering the role of E-cadherin and EMT in tumor spreading, it follows that PARP12-mediated MARylation of structural proteins such as Golgin-97 can be considered one of the central events in carcinogenesis.

## Materials and Methods

### Plasmids

The plasmids used in this study were generated by standard molecular cloning procedures as, described in details in the Appendix Method section.

### Antibodies

Antibodies were used as follows (final dilutions are indicated): PARP12-rabbit polyclonal antibody (Sigma-Prestige) was used for western blotting at 1:2000; PARP12-goat polyclonal antibody (Abcam) was used for immunofluorescence at 1:200; Golgin-97-rabbit polyclonal antibody (Sigma-Aldrich) was used for western blotting at 1:2000 and for immunofluorescence at 1:200; Golgin-245-mouse monoclonal antibody (BD Transduction Laboratories) was used at 1:1000 for western blotting and at 1:100 for immunofluorescence; tankyrase-1/2-rabbit polyclonal antibody (Santa Cruz Biotechnology) was used at 1:1000 for western blotting; TGN46-sheep polyclonal antibody (Bio-Rad) was used at 1:200 for immunofluorescence; GM130-mouse monoclonal antibody (BD Transduction Laboratories) was used at 1:100 for immunofluorescence; Giantin-mouse monoclonal antibody (Enzo Life Sciences) was used at 1:1000 for immunofluorescence; LAMP1-rabbit polyclonal antibody (Abcam) was used for immunofluorescence at 1:200; EEA1-mouse monoclonal antibody (BD Transduction Laboratories) was used at 1:100 for immunofluorescence; Transferrin Receptor/CD71 monoclonal antibody (clone H68.4, ThermoFisher Scientific) was used at 1:200 for immunofluorescence; FLAG-mouse monoclonal antibody (Sigma-Aldrich) was used at 1:5000 for western blotting and at 1:500 for immunofluorescence; GFP-mouse monoclonal (Abcam) was used at 1:100 for immunofluorescence. Sources of all antibodies and reagents used are listed in Table EV1.

### Cell Culture and treatments

HeLa cells (wild-type, ΔGolgin-97 or ΔGolgin-245 HeLa cells, kindly provided by Sean Munro, Shin et al., 2017) and MCF7 cells (from ATCC) were cultured in Dulbecco’s modified Eagle’s medium (DMEM) supplemented with 10% fetal bovine serum (FBS) and penicillin/streptomycin at 37 °C and 5% CO2. All cell culture reagents were from Life Technologies. For treatment with PJ34 (50 μM) or with IWR (25 μM), cells were grown to 80% confluence and then treated with inhibitors for 2 h. Nocodazole treatment (33 μM) was performed for 3 h. All treatments were carried out in complete growth media.

### Transient transfections

HeLa cells were transfected with plasmids encoding FLAG-tagged Golgin-97 (wild-type or its ADP-ribosylation-defective point mutants), mCherry-tagged Rab11, GFP-tagged E-cadherin or with constructs encoding E-cadherin and TNFα RUSH system using TransIT-LT1 Reagent (Mirus, MIR 2305) according to the manufacturer’s instructions.

### siRNA-mediated knockdown

Commercially available siRNA oligos targeting human PARP12 (Dharmacon, L-013740-00-0005), human Golgin-97 (synthetized by Sigma-Aldrich; #1: AAGAUCACAGCCCUGGAACAA[dT]; #2: AAGUGCUUCUCCAGAAAGAGC[dT] as reported by Lock et al., 2005) or non-targeting siRNAs (Dharmacon, D-001810-01-05) were transfected in HeLa and MCF7 cells at a final concentration of 100 or 50 nM using Lipofectamine RNAiMAX reagent (Invitrogen, 13778150) for 72 and 48 h, respectively, according to the manufacturer instructions. Where indicated, after 24 h, cells were transiently transfected with FLAG-tagged Golgin-97 or with constructs encoding E-cadherin and TNFα used for the RUSH system.

### Purification of His-tagged Golgin-97 from *Escherichia coli*

His-tagged Golgin-97 was purified as described below. The following procedure led to about 2 mg of recombinant His-tagged protein. BL21-DE3 cells (ThermoFisher Scientific, EC0114) transformed with the plasmid encoding the His-tagged Golgin-97 were scraped from the glycerol stocks, inoculated into 5 ml Luria broth (LB) containing 60 μg/ml kanamycin, and grown overnight at 37 °C under continuous shaking (200 rpm). The cultures were then diluted in 400 ml LB, and OD_600_ was monitored until 0.6. Bacteria were then induced with 0.2 mM IPTG at 25 °C. After 4 h, the cultures were chilled on ice and centrifuged at 5,000× *g* for 10 min at 4 °C. The pellets were resuspended in 4 ml His-lysis buffer (50 mM Tris-HCl, pH 8, 300 mM NaCl, 10% glycerol, 5 mM imidazole, 10 mM 2-mercaptoethanol, protease inhibitor cocktail), and frozen by immersion in liquid nitrogen. The lysates were stored at −80 °C overnight, or for a few days. Each suspension was thawed at 4 °C in a water bath, and both protease inhibitor cocktail and lysozyme (0.5 mg/ml) were added. Lysates were then incubated at 4 °C with gentle shaking. After 30 min, 1.5% Triton X-100 was added and the lysates incubated at 4 °C for further 30 min. Lysates were sonicated on ice three times for 30 sec and then centrifuged at 20,000× *g* for 20 min at 4 °C. The resulting supernatants were recovered and incubated with 0.5 ml Ni-NTA beads (Qiagen, 30230), previously equilibrated in His-lysis buffer. Each suspension was incubated with gentle shaking at 4 °C for 2 h, and then packed into a 10 ml chromatography column (Bio-Rad Laboratories, 7311550). The columns were washed 5 times with 10 ml His-wash buffer *plus* 20 mM imidazole, and the protein was eluted by adding aliquots of 0.5 ml His-elution buffer *plus* 300 mM imidazole, with each eluate collected in a clean tube. The elution and collection steps were repeated at least 5 times. The protein peaked in the first two fractions. The fractions containing the greater amounts of protein (at least 0.2 mg/ml) were pooled, dialyzed twice against 1,000 vol. of PBS *plus* 10% glycerol, flash frozen in liquid nitrogen, and stored in aliquots at −80 °C. The same procedure was used for the purification of wild-type Golgin-97 and Golgin-97 mutants (Mut A: E381Q-E386Q-E393Q; B: E558Q-E559Q-E565Q; C: E579Q-E585Q).

### Purification of GST-tagged Af1521 *macro* domain from Escherichia coli

The procedure to obtain the Af1521 *macro* domain resin was described previously (Grimaldi et al., 2018). BL21-DE3 cells (ThermoFisher Scientific, EC0114) transformed with the plasmid encoding the *macro* domain were scraped from the glycerol stocks, inoculated into 50 ml LB containing 100 μg/ml ampicillin, and grown overnight at 37 °C under continuous shaking (200 rpm). The cultures were then diluted 1:10 in 500 ml of the same medium, and the OD_600_ was monitored until it reached 0.6. The bacteria were then induced with the addition of 0.2 mM IPTG, for 3.5 h at 20 °C. After this, the cultures were chilled on ice and centrifuged at 5,000× g at 4 °C. The pellets were then resuspended in 10 ml STE buffer (150 mM NaCl, 20 mM Tris HCl, pH 8, 1 mM EDTA) containing protease inhibitor cocktail, 1 mM dithiothreitol and 0.5 mg/ml lysozyme, incubated with gentle shaking for 30 min at 4 °C and then frozen by immersion in liquid nitrogen and stored at −80 °C overnight, or for up to a few days. The suspension was thawed and 1% (w/v) Triton X-100, protease inhibitor cocktail and 1 mM DTT were added. The lysate was incubated with gentle agitation at 4 °C for 20 min, sonicated on ice three times for 30 sec and clarified by centrifugation at 20,000× g for 15 min at 4 °C. The resulting supernatant was incubated with 0.5 ml glutathione Sepharose matrix (previously equilibrated in washing buffer), for 1 h on a rotating wheel at 4 °C and then washed 6 times with 12 ml washing buffer (PBS, 1 mM EDTA, 1 mM dithiothreitol). The resin was then treated with the cross-linker DMP, as follows. First, the resin was washed twice with 10 ml 0.2 M sodium borate, pH 8.6, and centrifuged at 1,000× g for 5 min at room temperature. All of the further centrifugation steps were performed under the same conditions. Then, the suspension was incubated with 600 μl 0.2 M Tris-ethanolamine, pH 8.3, and 20 mM of the cross-linker DMP was added, with incubation on a rocker for 30 min at room temperature. At the end of this incubation, the suspension was centrifuged, and the supernatant removed. The reaction was stopped by washing the beads once with 14 ml 0.2 M ethanolamine, pH 8.2, and then incubating in the same buffer (10 ml) for 1 hour at room temperature on a rocker. At the end of this incubation, the beads were washed three times in PBS, three times with 20 mM GSH, in 20 mM Tris HCl, pH 8, once again in PBS, and finally with 100 mM glycine, pH 2.5. Finally, the cross-linked matrice was washed twice with PBS and stored in PBS with 0.02% (w/v) sodium azide at 4 °C. Alternatively, resins were not treated with the cross-linker DMP, but were washed 5 times with washing buffer and GST-tagged *macro* domain eluted in elution buffer (100 mM Tris, pH 8.0, 20 mM reduced glutathione, 5 mM DTT) and collected in a clean tube. The elution and collection steps were repeated 5 times. The protein peaked in the first three fractions. The fractions containing the greater amounts of protein were pooled, dialyzed twice against 1,000 vol. of PBS, flash frozen in liquid nitrogen, and stored in aliquots at −80 °C.

### *In Vitro* ADP-ribosylation assay

Purified GST-tagged PARP12 catalytic fragment (250 ng) was incubated with 1 μg purified recombinant wild-type Golgin-97 in ADP-ribosylation Buffer [50 mM Tris-HCl pH 7.4, 4 mM DTT, 500 μM MgCl_2_, 30 μM unlabeled NAD^+^ /4 μCi of ^32^P-NAD^+^] at 37 °C for 60 min. The ADP-ribosylation reactions were stopped by the addition of 2x SDS-PAGE loading buffer followed by heating to 95°C for 10 min. Samples were subjected to SDS-PAGE (8% PAGE-SDS gel) and transferred onto PVDF membranes. The incorporated [^32^P]-ADP-ribose was detected by autoradiography. Alternatively, purified GST-tagged PARP12 catalytic fragment (500 ng) was incubated with 3 μg purified recombinant Golgin-97 (wild-type and the different multiple point mutants) in ADP-ribosylation Buffer devoid of labeled NAD^+^ [50 mM Tris-HCl pH 7.4, 4 mM DTT, 500 μM MgCl_2_, 100 μM unlabeled NAD^+^] at 37 °C for 60 min. At the end of the reaction, 1/10 of the reaction mixture was saved (input material); 10 μg of purified GST-tagged Af1521 *macro* domain were added to the remaining reaction mix and incubated at 4 °C on a rotating wheel. After 2 h, samples were recovered, glutathione Sepharose matrix (50 μl of bed volume, previously equilibrated in reaction buffer) added to the samples and further incubated at 4 °C on a rotating wheel. After 1 h incubation, samples were recovered and washed three times in reaction buffer. Each time, the samples were centrifuged at 500× *g* for 5 min. SDS-PAGE loading buffer was added to the samples, followed by heating to 95°C for 5 min. Samples were subjected to SDS-PAGE (8% PAGE-SDS gel) and transferred onto nitrocellulose membranes for Western blotting using anti-Golgin-97 antibody. The signals obtained were acquired and quantified using a Gel-Doc system (Bio-Rad). The percentage of MARylated Golgin-97 was calculated as ratio between Af1521-bound Golgin-97/total Golgin-97.

### Macro domain based pull-down assay

HeLa cells were washed three times in ice-cold PBS and solubilized in RIPA buffer (100 mM Tris HCl, pH 7.5, 150 mM NaCl, 1% NP-40, 0.5% deoxycholate, 0.1% SDS, supplemented with protease inhibitors and 5 μM PJ34), under constant rotation for 30 min at 4 °C. The mixtures were clarified by centrifugation at 13,000× *g* for 10 min at 4 °C, the supernatants were recovered and protein concentration evaluated using the BCA Protein Assay Kit (Pierce, 23225). Total lysates (1 mg) were then incubated overnight with 50 μl of a 10 μg/μl GST cross-linked wild-type macro-domain resin at 4 °C, on a rotating wheel. After the incubation, the mixtures were centrifuged at 500× *g* for 5 min to recover the proteins bound to the wild-type macro domain. The resin was previously equilibrated with RIPA buffer. Resins were then washed 3 times with RIPA buffer and another 2 times in the same buffer without detergents. Each time, the samples were centrifuged at 500× *g* for 5 min. At the end of the washing steps, the resins were resuspended in 100 μl SDS sample buffer, boiled, analyzed by 8% SDS/PAGE and transferred onto nitrocellulose for Western blotting.

The membranes were blocked for 1 h at room temperature in Tris-Buffered Saline (TBS) with 0.05% Tween (TBS-T) containing 5% BSA. Primary antibodies were diluted (see “Antibodies” section above for specific dilutions used) in blocking solution and incubated with membranes overnight at 4° C with gentle mixing. After extensive washing with TBS-T, the membranes were incubated with an appropriate HRP-conjugated secondary antibody (Millipore) diluted in 5% non-fat, dry milk in TBS-T for 45 min at room temperature. The membranes were washed extensively with TBS-T before chemiluminescent detection using the ECL western Blotting Detection Reagents (GE Healthcare Life Sciences, RPN2106) and X-ray film or VersaDoc system (Bio-Rad).

### Immunofluorescent Staining and Confocal Microscopy

Cells were fixed in 4% paraformaldehyde for 10 min at room temperature, washed three times in PBS, and incubated for 30 min at room temperature in blocking solution (0.5% bovine serum albumin, 50 nM NH_4_Cl in PBS, pH 7.4, 0.1% saponin and 0.02% sodium azide). The cells were subsequently incubated with the indicated antibodies diluted in blocking solution (see above for dilutions used) for 2 h at room temperature or overnight at 4 °C. After incubation with the primary antibody, cells were washed three times in PBS and incubated with a fluorescent-probe-conjugated secondary antibody, for 30 min at room temperature. Alexa Fluor 488-, 568- or 647-conjugated anti-rabbit, anti-mouse or anti-goat donkey antibodies were used at a dilution of 1:400 in blocking solution. After immuno-staining, cells were washed three times in PBS and twice in sterile water, to remove salts. The coverslips were then mounted on glass-microscope slides with Mowiol (20 mg mowiol dissolved in 80 ml PBS, stirred overnight and centrifuged for 30 min at 12,000× *g*). Images were taken using a Zeiss-LSM 700 confocal microscope. Optical confocal sections were taken at 1 Air Unit.

### Transport assays

#### TGN-exit assay of VSVG

HeLa cells (wild-type and ΔGolgin-97) were infected with vesicular stomatitis virus (VSV) and incubated overnight at 40 °C, following incubation at 20 °C for 90 min in presence of cycloheximide (50 μg/ml) to accumulate the G protein of VSV (VSVG) in the Golgi complex. To monitor VSVG exit from the TGN the temperature was shifted to 32 °C and the samples were fixed with 4% paraformaldehyde at different times (40° and 20° C temperature blocks; 15, 30, 45 and 60 min of the 32° C temperature release) and stained with the indicated antibodies, as reported. Quantification was performed by counting the number of cells showing VSVG at the Golgi complex.

#### RUSH system

HeLa cells were plated on coverslips in 24-well plates and transfected with the Retention Using a Selective Hook (RUSH)-based EGFP-tagged TNFα or E-cadherin constructs (Str-KDEL_SBP-EGFP-E-cadherin/TNFα). After 24- or 48-h incubation at 37 °C, respectively, biotin (40 μM) was added at different times (0, 30, 60, 120 min) to allow the cargo to undergo trafficking. Cells were then fixed and stained with the specific antibodies, as reported. For surface immuno-staining, cells were fixed with freshly-prepared 4% paraformaldehyde and incubated with blocking solution without saponin. After 30 min, non-permeabilized cells were incubated with an anti-GFP antibody for 1 h at room temperature, washed three times with PBS and then incubated for further 30 min with the Alexa-568 secondary antibody. Samples were then analyzed using a Zeiss-LSM 700 confocal microscope.

#### Live cell imaging

For live cell imaging, wild-type or ΔGolgin-97 HeLa cells were grown on glass-bottomed 35-mm dishes and transiently transfected with GFP-tagged E-cadherin alone or GFP-tagged E-cadherin/Tomato-tagged Golgin-97 (wild-type, Mut A, Mut B, Mut C). Three hours after transfection, the glass-bottomed dishes were mounted on a Zeiss LSM700 laser scanning microscope under controlled temperature and CO_2_; cells showing GFP-E-cadherin at the Golgi complex were imaged for the duration of the experiment (2 h). Frames were taken every 3 min (488 for excitation; PMT: 510–550 nm; 512×512 pixels; frame average, 4).

### Quantitative fluorescence image analysis

Quantitative analysis was performed using the Image J software. In brief, to calculate the amount of cargo in the Golgi after a traffic pulse, the integrated intensity fluorescence was measured for each cell area and for the Golgi area of that cell and the Golgi/Total ratio was calculated. For quantification of the GFP-surface staining (used for the RUSH system), the integrated intensity fluorescence was measured for each cell area and for the cell surface and the Surface/Total ratio was calculated.

### Statistical Analysis

p values were calculated comparing control and each treated group individually using Student’s t test. All statistical parameters are listed in the corresponding figure legends.

## ACKNOWLEDGEMENTS

We thank Drs. S. Munro and J.J.H. Shin (MRC Laboratory of Molecular Biology, Cambridge, UK) for kindly providing the ΔGolgin-97, ΔGolgin-245 and the parental cell lines; Dr. P. Gleeson (University of Melbourne, Australia) for providing the Myc-tagged Golgin-245 construct; Dr. J. Stow (University of Queensland, Brisbane, Australia) for the GFP-tagged E-cadherin construct; Dr. B. Goud (Institut Curie, Paris, France) for providing mCherry-tagged Rab11 construct; Dr. M.A. De Matteis (Telethon Institute of Genetics and Medicine, Pozzuoli, Italy) for providing the FLAG-tagged wild-type Golgin-97 construct. Drs. A. Luini and R. Parashuraman (Institute of Biochemistry and Cell Biology, IBBC, CNR) for discussion and critical reading of the manuscript; Dr. G. Catara for help with cloning and construct preparation and Dr. Angela Filograna for contributing on MCF7 cell line work; the BioImaging Facility at the IBBC for support in imaging microscopy, data processing and analysis; the Italian Association for Cancer Research (AIRC) (to D.C. IG18776), the AIRC-Fondazione Cariplo TRansforming IDEas in Oncological research project (TRIDEO) (to C.V. IG17524), the PRONAT, PRIN No 20177XJCHX projects, the SATIN and CIRO POR-projects 2014-2020 for supporting our work.

## AUTHOR CONTRIBUTIONS

G.G. and D.C. designed the project and wrote the manuscript; G.G. and L.S. performed biochemical assays; M.L.M. and A.R.B. carried-out the bioinformatics analyses and residue identification; D.S. generated constructs; G.G., R.D.M. and C.V. performed transport assays and morphological analyses by confocal microscopy; all authors regularly discussed approaches and data.

## Expanded View Information

### Expanded View Figure Legends

**Fig EV1. E-cadherin-transport carriers colocalize with Rab11**

ΔGolgin-97 HeLa cells were transiently transfected with GFP-tagged E-cadherin (green). After 24 h overexpression, cells were fixed and stained with markers of the Golgi apparatus (GM130, Giantin, TGN46), early endosomes (EEA1), lysosomes (LAMP1) and recycling endosomes (Transferrin receptor; TnfR), as indicated. **(B)** To follow Rab11 localization, ΔGolgin-97 HeLa cells were co-transfected with mCherry-tagged Rab11 and GFP-tagged E-cadherin; after 12 h transfection cells were fixed and processed for immunofluorescence analysis. Insets: enlarged view of merged signals. Scale bars, 10 μm. Note that the pool of E-cadherin at the Golgi compartment did not colocalize with GM130 or Giantin, excluding the cargo from the cis/medial side of the Golgi complex (intra-Golgi defect); instead, it clearly co-localized with TGN46 (A). Of note, the E-cadherin-including carriers did not show any degree of colocalization with either EEA1 or LAMP1 (A), while they colocalized with Rab11 (B), defining a post-TGN, Rab11-positive intermediate compartment as the site of E-cadherin accumulation.

In addition, the E-cadherin-containing carriers did not co-localize with TnfR (while they co-localized with Rab11). This suggests that these carriers are not recycling vesicles, but define a post-TGN, Rab11-positive intermediate compartment.

**Fig EV2. VSVG transport is inhibited in the absence of Golgin-97**

Wild-type (**A**) and ΔGolgin-97 (**B**) HeLa cells were infected with VSV virus (see methods). After the 20 °C temperature-block, cells were incubated at 32 °C to allow the release of the cargo from the TGN, fixed at different times and stained with VSVG (green), TGN46 and Golgin-97 (cyan) antibodies. Representative confocal images of cells fixed at 30 min are shown. (**C**) Quantification of cells showing VSVG in the Golgi area after 30 min of the 32 °C temperature-block release is reported. Data are mean of n= 300 cells from three different experiments ± SD. Statistical significance was calculated by unpaired Student’s t-test; ***p < 0.001 Scale bars, 10 μm.

**Fig EV3. ADP-ribosylation site prediction for Golgin-97**

Graphic visualization of predicted ADP-ribosylation sites for Golgin-97, as downloaded from ADPredict web application (www.adpredict.net). Histogram bars report the total score for each acidic residue, the higher the most prone to be modified.

**Fig EV4. E-cadherin transport in ΔGolgin-97 cells following expression of wild-type Golgin-97 or its ADP-ribosylation defective mutant**

Representative confocal images of Movies EV3 and EV4 for GFP-tagged E-cadherin (gray) transport in ΔGolgin-97 HeLa cells transiently transfected with the Tomato-tagged wild-type Golgin-97 or with its ADP-ribosylation defective mutant B (Mut B, red), as indicated. Scale bars, 10 μm.

**Fig EV5. Absence of Golgin-97 alters endogenous E-cadherin localization**

MCF7 cells were transiently transfected with Golgin-97 siRNAs. After 48 h, cells were fixed and stained with anti-E-cadherin (green) and Golgin-97 (cyan) antibodies. Representative confocal images are shown. Scale bars, 10 μm. Note that the absence of Golgin-97 impairs the E-cadherin localization at the PM.

## Expanded View Movies

### Movie EV1

Transport of GFP-E-cadherin in wild-type HeLa cells, related to Fig 5. Live cell confocal imaging of GFP-E-cadherin transfected wild-type HeLa cells. Three hours after transfection, cells showing GFP-E-cadherin (gray) at the Golgi complex were imaged for the duration of the experiment (2 h). Frames were taken every 3 min.

### Movie EV2

Transport of GFP-E-cadherin in ΔGolgin-97 HeLa cells, related to Figure 5. Live cell confocal imaging of GFP-E-cadherin transfected ΔGolgin-97 HeLa cells. Three hours after transfection, cells showing GFP-E-cadherin (gray) at the Golgi complex were imaged for the duration of the experiment (2 h). Frames were taken every 3 min.

### Movie EV3

Transport of GFP-E-cadherin in ΔGolgin-97 HeLa cells, co-transfected with Tomato-wild-type Golgin-97, related to Fig 5. Three hours after transfection, cells showing GFP-E-cadherin at the Golgi complex (gray) were imaged for the duration of the experiment (2 h). Frames were taken every 3 min.

### Movie EV4

Transport of GFP-E-cadherin in ΔGolgin-97 HeLa cells, co-transfected with Tomato-Mut B Golgin-97, related to Fig 5. Three hours after transfection, cells showing GFP-E-cadherin at the Golgi complex (gray) were imaged for the duration of the experiment (2 h). Frames were taken every 3 min.

**Table EV1:**
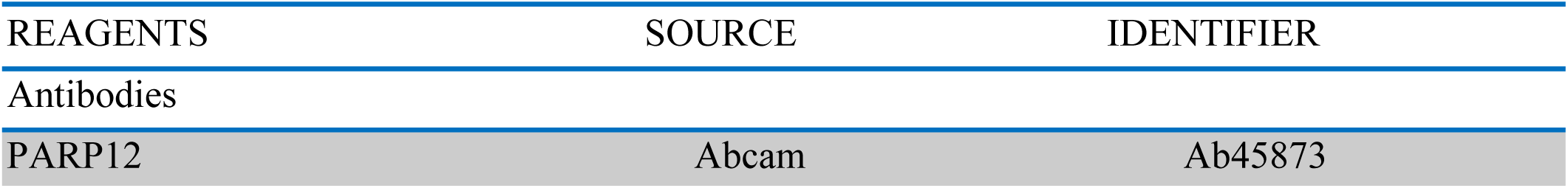
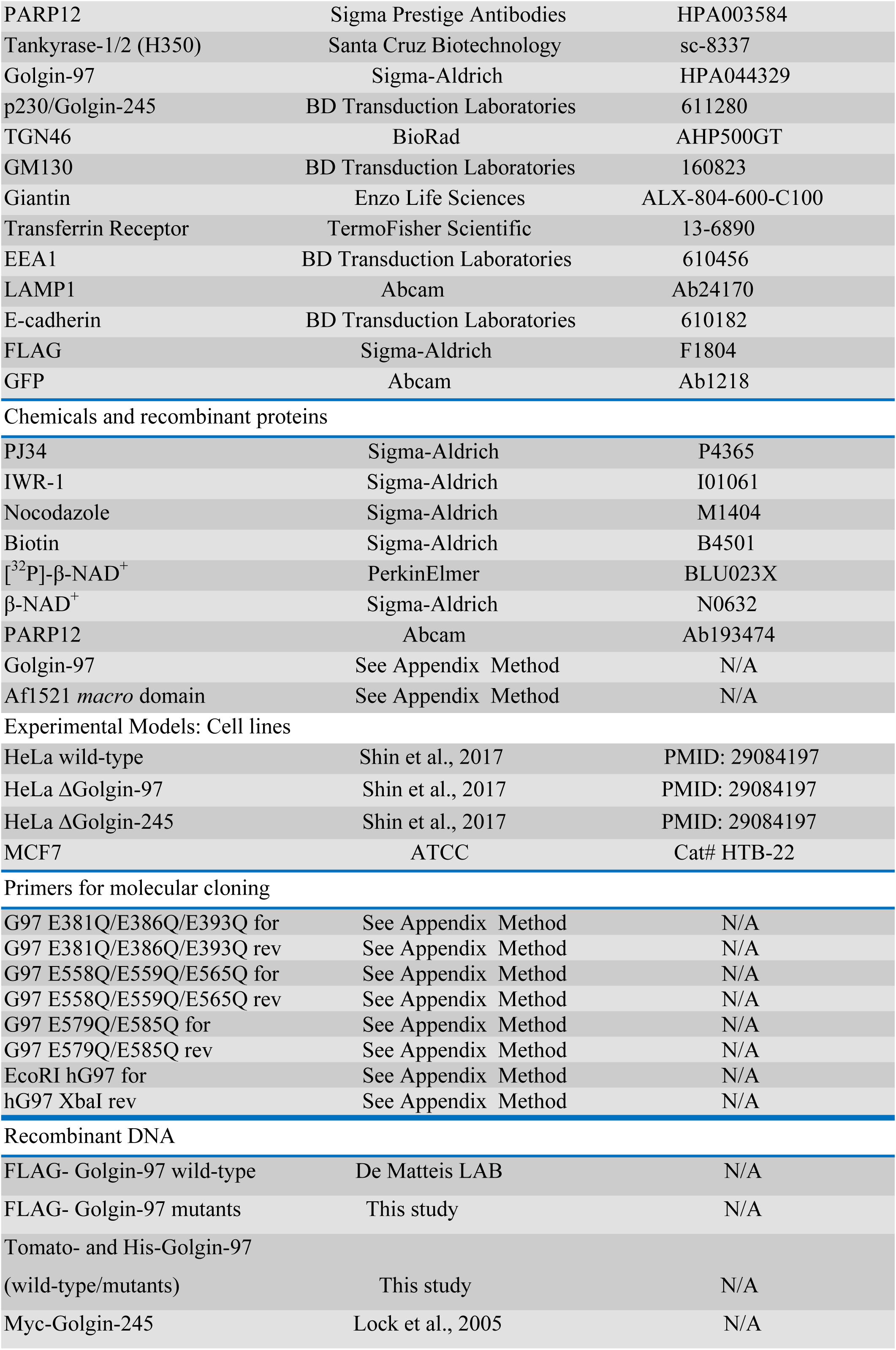
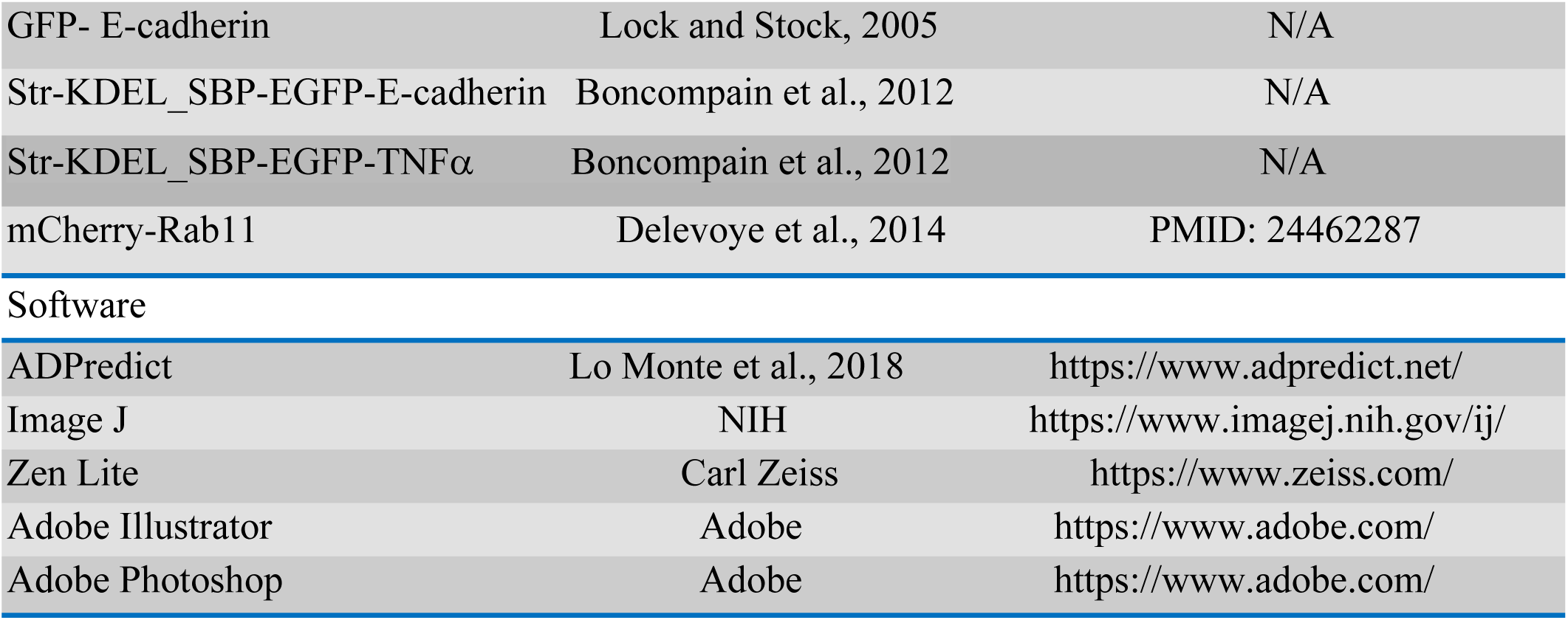
RESOURCES TABLE.

| REAGENTS | SOURCE | IDENTIFIER |
| --- | --- | --- |
| Antibodies |  |  |
| PARP12 | Abcam | Ab45873 |
| PARP12 | Sigma Prestige Antibodies | HPA003584 |
| Tankyrase-1/2 (H350) | Santa Cruz Biotechnology | sc-8337 |
| Golgin-97 | Sigma-Aldrich | HPA044329 |
| p230/Golgin-245 | BD Transduction Laboratories | 611280 |
| TGN46 | BioRad | AHP500GT |
| GM130 | BD Transduction Laboratories | 160823 |
| Giantin | Enzo Life Sciences | ALX-804-600-C100 |
| Transferrin Receptor | TermoFisher Scientific | 13-6890 |
| EEA1 | BD Transduction Laboratories | 610456 |
| LAMP1 | Abcam | Ab24170 |
| E-cadherin | BD Transduction Laboratories | 610182 |
| FLAG | Sigma-Aldrich | F1804 |
| GFP | Abcam | Ab1218 |
| Chemicals and recombinant proteins |  |  |
| PJ34 | Sigma-Aldrich | P4365 |
| IWR-1 | Sigma-Aldrich | I01061 |
| Nocodazole | Sigma-Aldrich | M1404 |
| Biotin | Sigma-Aldrich | B4501 |
| [<sup>32</sup>P]-β-NAD<sup>+</sup> | PerkinElmer | BLU023X |
| β-NAD<sup>+</sup> | Sigma-Aldrich | N0632 |
| PARP12 | Abcam | Ab193474 |
| Golgin-97 | See Appendix Method | N/A |
| Af1521 *macro* domain | See Appendix Method | N/A |
| Experimental Models: Cell lines |  |  |
| HeLa wild-type | Shin et al., 2017 | PMID: 29084197 |
| HeLa ΔGolgin-97 | Shin et al., 2017 | PMID: 29084197 |
| HeLa ΔGolgin-245 | Shin et al., 2017 | PMID: 29084197 |
| MCF7 | ATCC | Cat# HTB-22 |
| Primers for molecular cloning |  |  |
| G97 E381Q/E386Q/E393Q for | See Appendix Method | N/A |
| G97 E381Q/E386Q/E393Q rev | See Appendix Method | N/A |
| G97 E558Q/E559Q/E565Q for | See Appendix Method | N/A |
| G97 E558Q/E559Q/E565Q rev | See Appendix Method | N/A |
| G97 E579Q/E585Q for | See Appendix Method | N/A |
| G97 E579Q/E585Q rev | See Appendix Method | N/A |
| EcoRI hG97 for | See Appendix Method | N/A |
| hG97 XbaI rev | See Appendix Method | N/A |
| Recombinant DNA |  |  |
| FLAG- Golgin-97 wild-type | De Matteis LAB | N/A |
| FLAG- Golgin-97 mutants | This study | N/A |
| Tomato- and His-Golgin-97<br>(wild-type/mutants) | This study | N/A |
| Myc-Golgin-245 | Lock et al., 2005 | N/A |
| GFP- E-cadherin | Lock and Stock, 2005 | N/A |
| Str-KDEL_SBP-EGFP-E-cadherin | Boncompain et al., 2012 | N/A |
| Str-KDEL_SBP-EGFP-TNFα | Boncompain et al., 2012 | N/A |
| mCherry-Rab11 | Delevoye et al., 2014 | PMID: 24462287 |
| Software |  |  |
| ADPredict | Lo Monte et al., 2018 | https://www.adpredict.net/ |
| Image J | NIH | https://www.imagej.nih.gov/ij/ |
| Zen Lite | Carl Zeiss | https://www.zeiss.com/ |
| Adobe Illustrator | Adobe | https://www.adobe.com/ |
| Adobe Photoshop | Adobe | https://www.adobe.com/ |

## References

Abplanalp, J., Leutert, M., Frugier, E., Nowak, K., Feurer, R., Kato, J., Kistemaker, H.V.A., Filippov, D.V., Moss, J., Caflisch, A., et al. (2017). Proteomic analyses identify ARH3 as a serine mono-ADP-ribosylhydrolase. Nat Commun 8, 2055.

Atasheva, S., Akhrymuk, M., Frolova, E.I., and Frolov, I. (2012). New PARP gene with an anti-alphavirus function. J Virol 86, 8147–8160.

Atasheva, S., Frolova, E.I., and Frolov, I. (2014). Interferon-Stimulated Poly(ADP-Ribose) Polymerases Are Potent Inhibitors of Cellular Translation and Virus Replication. J Virol 88, 2116–2130.

Beznoussenko, G.V., Parashuraman, S., Rizzo, R., Polishchuk, R., Martella, O., Di Giandomenico, D., Fusella, A., Spaar, A., Sallese, M., Capestrano, M.G., et al. (2014). Transport of soluble proteins through the Golgi occurs by diffusion via continuities across cisternae. Elife 3.

Boncompain, G., Divoux, S., Gareil, N., de Forges, H., Lescure, A., Latreche, L., Mercanti, V., Jollivet, F., Raposo, G., and Perez, F. (2012). Synchronization of secretory protein traffic in populations of cells. Nat Methods 9, 493–498.

Bonfiglio, J.J., Fontana, P., Zhang, Q., Colby, T., Gibbs-Seymour, I., Atanassov, I., Bartlett, E., Zaja, R., Ahel, I., and Matic, I. (2017). Serine ADP-Ribosylation Depends on HPF1. Mol Cell 65, 932–940 e936.

Bremnes, R.M., Veve, R., Hirsch, F.R., and Franklin, W.A. (2002). The E-cadherin cell-cell adhesion complex and lung cancer invasion, metastasis, and prognosis. Lung Cancer 36, 115–124.

Bruser, L., and Bogdan, S. (2017). Adherens Junctions on the Move-Membrane Trafficking of E-Cadherin. Cold Spring Harb Perspect Biol 9.

Buchan, J.R., and Parker, R. (2009). Eukaryotic stress granules: the ins and outs of translation. Mol Cell 36, 932–941.

Cadwell, C.M., Su, W., and Kowalczyk, A.P. (2016). Cadherin tales: Regulation of cadherin function by endocytic membrane trafficking. Traffic 17, 1262–1271.

Catara, G., Grimaldi, G., Schembri, L., Spano, D., Turacchio, G., Lo Monte, M., Beccari, A.R., Valente, C., and Corda, D. (2017). PARP1-produced poly-ADP-ribose causes the PARP12 translocation to stress granules and impairment of Golgi complex functions. Sci Rep 7, 14035.

Chang, P., Coughlin, M., and Mitchison, T.J. (2005a). Tankyrase-1 polymerization of poly(ADP-ribose) is required for spindle structure and function. Nat Cell Biol 7, 1133–1139.

Chang, P., Coughlin, M., and Mitchison, T.J. (2009). Interaction between Poly(ADP-ribose) and NuMA contributes to mitotic spindle pole assembly. Mol Biol Cell 20, 4575–4585.

Chang, W., Dynek, J.N., and Smith, S. (2005b). NuMA is a major acceptor of poly(ADP-ribosyl)ation by tankyrase 1 in mitosis. Biochem J 391, 177–184.

Chen, H.C., Chu, R.Y., Hsu, P.N., Hsu, P.I., Lu, J.Y., Lai, K.H., Tseng, H.H., Chou, N.H., Huang, M.S., Tseng, C.J., et al. (2003). Loss of E-cadherin expression correlates with poor differentiation and invasion into adjacent organs in gastric adenocarcinomas. Cancer Lett 201, 97–106.

Chen, Y., Gershlick, D.C., Park, S.Y., and Bonifacino, J.S. (2017). Segregation in the Golgi complex precedes export of endolysosomal proteins in distinct transport carriers. J Cell Biol 216, 4141–4151.

Cheung, P.Y., and Pfeffer, S.R. (2016). Transport Vesicle Tethering at the Trans Golgi Network: Coiled Coil Proteins in Action. Front Cell Dev Biol 4, 18.

Chi, N.W., and Lodish, H.F. (2000). Tankyrase is a golgi-associated mitogen-activated protein kinase substrate that interacts with IRAP in GLUT4 vesicles. J Biol Chem 275, 38437–38444.

Cohen, M.S., and Chang, P. (2018). Insights into the biogenesis, function, and regulation of ADP-ribosylation. Nat Chem Biol 14, 236–243.

Corda, D., and Di Girolamo, M. (2003). Functional aspects of protein mono-ADP-ribosylation. EMBO J 22, 1953–1958.

Dani, N., Mayo, E., Stilla, A., Marchegiani, A., Di Paola, S., Corda, D., and Di Girolamo, M. (2011). Mono-ADP-ribosylation of the G protein betagamma dimer is modulated by hormones and inhibited by Arf6. J Biol Chem 286, 5995–6005.

Dani, N., Stilla, A., Marchegiani, A., Tamburro, A., Till, S., Ladurner, A.G., Corda, D., and Di Girolamo, M. (2009). Combining affinity purification by ADP-ribose-binding macro domains with mass spectrometry to define the mammalian ADP-ribosyl proteome. Proc Natl Acad Sci U S A 106, 4243–4248.

De Matteis, M.A., and Luini, A. (2008). Exiting the Golgi complex. Nat Rev Mol Cell Biol 9, 273–284.

Di Girolamo, M., Dani, N., Stilla, A., and Corda, D. (2005). Physiological relevance of the endogenous mono(ADP-ribosyl)ation of cellular proteins. FEBS J 272, 4565–4575.

Di Paola, S., Micaroni, M., Di Tullio, G., Buccione, R., and Di Girolamo, M. (2012). PARP16/ARTD15 is a novel endoplasmic-reticulum-associated mono-ADP-ribosyltransferase that interacts with, and modifies karyopherin-ss1. PLoS One 7, e37352.

Dolle, C., Niere, M., Lohndal, E., and Ziegler, M. (2010). Visualization of subcellular NAD pools and intra-organellar protein localization by poly-ADP-ribose formation. Cell Mol Life Sci 67, 433–443.

Fan, X., Jin, S., Li, Y., Khadaroo, P.A., Dai, Y., He, L., Zhou, D., and Lin, H. (2019). Genetic And Epigenetic Regulation Of E-Cadherin Signaling In Human Hepatocellular Carcinoma. Cancer Manag Res 11, 8947–8963.

Fontana, P., Bonfiglio, J.J., Palazzo, L., Bartlett, E., Matic, I., and Ahel, I. (2017). Serine ADP-ribosylation reversal by the hydrolase ARH3. Elife 6.

Gill, D.M., Pappenheimer, A.M., Jr., Brown, R., and Kurnick, J.T. (1969). Studies on the mode of action of diphtheria toxin. VII. Toxin-stimulated hydrolysis of nicotinamide adenine dinucleotide in mammalian cell extracts. J Exp Med 129, 1–21.

Grimaldi, G., Catara, G., Palazzo, L., Corteggio, A., Valente, C., and Corda, D. (2019). PARPs and PAR as novel pharmacological targets for the treatment of stress granule-associated disorders. Biochem Pharmacol.

Grimaldi, G., Catara, G., Valente, C., and Corda, D. (2018). In Vitro Techniques for ADP-Ribosylated Substrate Identification. Methods Mol Biol 1813, 25–40.

Grimaldi, G., and Corda, D. (2019). ADP-ribosylation and intracellular traffic: an emerging role for PARP enzymes. Biochem Soc Trans 47, 357–370.

Guo, H.L., Zhang, C., Liu, Q., Li, Q., Lian, G., Wu, D., Li, X., Zhang, W., Shen, Y., Ye, Z., et al. (2012). The Axin/TNKS complex interacts with KIF3A and is required for insulin-stimulated GLUT4 translocation. Cell Res 22, 1246–1257.

Gupte, R., Liu, Z., and Kraus, W.L. (2017). PARPs and ADP-ribosylation: recent advances linking molecular functions to biological outcomes. Genes Dev 31, 101–126.

Hanahan, D., and Weinberg, R.A. (2011). Hallmarks of cancer: the next generation. Cell 144, 646–674.

Hassa, P.O., and Hottiger, M.O. (2008). The diverse biological roles of mammalian PARPS, a small but powerful family of poly-ADP-ribose polymerases. Front Biosci 13, 3046–3082.

Hottiger, M.O., Hassa, P.O., Luscher, B., Schuler, H., and Koch-Nolte, F. (2010). Toward a unified nomenclature for mammalian ADP-ribosyltransferases. Trends Biochem Sci 35, 208–219.

Jing, J., Junutula, J.R., Wu, C., Burden, J., Matern, H., Peden, A.A., and Prekeris, R. (2010). FIP1/RCP binding to Golgin-97 regulates retrograde transport from recycling endosomes to the trans-Golgi network. Mol Biol Cell 21, 3041–3053.

Jones, E.M., and Baird, A. (1997). Cell-surface ADP-ribosylation of fibroblast growth factor-2 by an arginine-specific ADP-ribosyltransferase. Biochem J 323 (Pt 1), 173–177.

Jwa, M., and Chang, P. (2012). PARP16 is a tail-anchored endoplasmic reticulum protein required for the PERK- and IRE1alpha-mediated unfolded protein response. Nat Cell Biol 14, 1223–1230.

Kedersha, N., and Anderson, P. (2002). Stress granules: sites of mRNA triage that regulate mRNA stability and translatability. Biochem Soc Trans 30, 963–969.

Kulkarni-Gosavi, P., Makhoul, C., and Gleeson, P.A. (2019). Form and function of the Golgi apparatus: scaffolds, cytoskeleton and signalling. FEBS Lett 593, 2289–2305.

Larsen, S.C., Hendriks, I.A., Lyon, D., Jensen, L.J., and Nielsen, M.L. (2018). Systems-wide Analysis of Serine ADP-Ribosylation Reveals Widespread Occurrence and Site-Specific Overlap with Phosphorylation. Cell Rep 24, 2493–2505 e2494.

Leung, A.K. (2014). Poly(ADP-ribose): an organizer of cellular architecture. J Cell Biol 205, 613–619.

Leung, A.K., Vyas, S., Rood, J.E., Bhutkar, A., Sharp, P.A., and Chang, P. (2011). Poly(ADP-ribose) regulates stress responses and microRNA activity in the cytoplasm. Mol Cell 42, 489–499.

Li, L., Zhao, H., Liu, P., Li, C., Quanquin, N., Ji, X., Sun, N., Du, P., Qin, C.F., Lu, N., et al. (2018). PARP12 suppresses Zika virus infection through PARP-dependent degradation of NS1 and NS3 viral proteins. Sci Signal 11.

Lieu, Z.Z., Lock, J.G., Hammond, L.A., La Gruta, N.L., Stow, J.L., and Gleeson, P.A. (2008). A trans-Golgi network golgin is required for the regulated secretion of TNF in activated macrophages in vivo. Proc Natl Acad Sci U S A 105, 3351–3356.

Lo Monte, M., Manelfi, C., Gemei, M., Corda, D., and Beccari, A.R. (2018). ADPredict: ADP-ribosylation site prediction based on physicochemical and structural descriptors. Bioinformatics 34, 2566–2574.

Lock, J.G., Hammond, L.A., Houghton, F., Gleeson, P.A., and Stow, J.L. (2005). E-cadherin transport from the trans-Golgi network in tubulovesicular carriers is selectively regulated by golgin-97. Traffic 6, 1142–1156.

Lowe, M. (2019). The Physiological Functions of the Golgin Vesicle Tethering Proteins. Front Cell Dev Biol 7, 94.

Lowe, M., Rabouille, C., Nakamura, N., Watson, R., Jackman, M., Jamsa, E., Rahman, D., Pappin, D.J., and Warren, G. (1998). Cdc2 kinase directly phosphorylates the cis-Golgi matrix protein GM130 and is required for Golgi fragmentation in mitosis. Cell 94, 783–793.

Luo, X., and Kraus, W.L. (2012). On PAR with PARP: cellular stress signaling through poly(ADP-ribose) and PARP-1. Genes Dev 26, 417–432.

Lupi, R., Corda, D., and Di Girolamo, M. (2000). Endogenous ADP-ribosylation of the G protein beta subunit prevents the inhibition of type 1 adenylyl cyclase. J Biol Chem 275, 9418–9424.

Lupi, R., Dani, N., Dietrich, A., Marchegiani, A., Turacchio, S., Berrie, C.P., Moss, J., Gierschik, P., Corda, D., and Di Girolamo, M. (2002). Endogenous mono-ADP-ribosylation of the free Gbetagamma prevents stimulation of phosphoinositide 3-kinase-gamma and phospholipase C-beta2 and is activated by G-protein-coupled receptors. Biochem J 367, 825–832.

Mann, M., and Jensen, O.N. (2003). Proteomic analysis of post-translational modifications. Nat Biotechnol 21, 255–261.

Moss, J., Balducci, E., Cavanaugh, E., Kim, H.J., Konczalik, P., Lesma, E.A., Okazaki, I.J., Park, M., Shoemaker, M., Stevens, L.A., et al. (1999). Characterization of NAD:arginine ADP-ribosyltransferases. Mol Cell Biochem 193, 109–113.

Moss, J., and Vaughan, M. (1977). Mechanism of action of choleragen. Evidence for ADP-ribosyltransferase activity with arginine as an acceptor. J Biol Chem 252, 2455–2457.

Muschalik, N., and Munro, S. (2018). Golgins. Curr Biol 28, R374–R376.

Nakamura, N., Lowe, M., Levine, T.P., Rabouille, C., and Warren, G. (1997). The vesicle docking protein p115 binds GM130, a cis-Golgi matrix protein, in a mitotically regulated manner. Cell 89, 445–455.

Okazaki, I.J., and Moss, J. (1996). Mono-ADP-ribosylation: a reversible posttranslational modification of proteins. Adv Pharmacol 35, 247–280.

Otto, H., Reche, P.A., Bazan, F., Dittmar, K., Haag, F., and Koch-Nolte, F. (2005). In silico characterization of the family of PARP-like poly(ADP-ribosyl)transferases (pARTs). BMC Genomics 6, 139.

Pagliuso, A., Valente, C., Giordano, L.L., Filograna, A., Li, G., Circolo, D., Turacchio, G., Marzullo, V.M., Mandrich, L., Zhukovsky, M.A., et al. (2016). Golgi membrane fission requires the CtBP1-S/BARS-induced activation of lysophosphatidic acid acyltransferase delta. Nat Commun 7, 12148.

Palacios, F., Tushir, J.S., Fujita, Y., and D’Souza-Schorey, C. (2005). Lysosomal targeting of E-cadherin: a unique mechanism for the down-regulation of cell-cell adhesion during epithelial to mesenchymal transitions. Mol Cell Biol 25, 389–402.

Palazzo, L., Leidecker, O., Prokhorova, E., Dauben, H., Matic, I., and Ahel, I. (2018). Serine is the major residue for ADP-ribosylation upon DNA damage. Elife 7.

Polishchuk, E.V., Di Pentima, A., Luini, A., and Polishchuk, R.S. (2003). Mechanism of constitutive export from the golgi: bulk flow via the formation, protrusion, and en bloc cleavage of large trans-golgi network tubular domains. Mol Biol Cell 14, 4470–4485.

Prieto-Garcia, E., Diaz-Garcia, C.V., Garcia-Ruiz, I., and Agullo-Ortuno, M.T. (2017). Epithelial-to-mesenchymal transition in tumor progression. Med Oncol 34, 122.

Seman, M., Adriouch, S., Haag, F., and Koch-Nolte, F. (2004). Ecto-ADP-ribosyltransferases (ARTs): emerging actors in cell communication and signaling. Curr Med Chem 11, 857–872.

Shin, J.J.H., Crook, O.M., Borgeaud, A., Cattin-Ortolá, J., Peak-Chew, S.-Y., Chadwick, J., Lilley, K.S., and Munro, S. (2019). Determining the content of vesicles captured by golgin tethers using LOPIT-DC. bioRxiv, 841965.

Shin, J.J.H., Gillingham, A.K., Begum, F., Chadwick, J., and Munro, S. (2017). TBC1D23 is a bridging factor for endosomal vesicle capture by golgins at the trans-Golgi. Nat Cell Biol 19, 1424–1432.

Slade, D. (2019). Mitotic functions of poly(ADP-ribose) polymerases. Biochem Pharmacol.

Sommariva, M., and Gagliano, N. (2020). E-Cadherin in Pancreatic Ductal Adenocarcinoma: A Multifaceted Actor during EMT. Cells 9.

Su, Z., Deshpande, V., James, D.E., and Stockli, J. (2018). Tankyrase modulates insulin sensitivity in skeletal muscle cells by regulating the stability of GLUT4 vesicle proteins. J Biol Chem 293, 8578–8587.

Valente, C., Turacchio, G., Mariggio, S., Pagliuso, A., Gaibisso, R., Di Tullio, G., Santoro, M., Formiggini, F., Spano, S., Piccini, D., et al. (2012). A 14-3-3gamma dimer-based scaffold bridges CtBP1-S/BARS to PI(4)KIIIbeta to regulate post-Golgi carrier formation. Nat Cell Biol 14, 343–354.

Vyas, S., Matic, I., Uchima, L., Rood, J., Zaja, R., Hay, R.T., Ahel, I., and Chang, P. (2014). Family-wide analysis of poly(ADP-ribose) polymerase activity. Nat Commun 5, 4426.

Welsby, I., Hutin, D., Gueydan, C., Kruys, V., Rongvaux, A., and Leo, O. (2014). PARP12, an interferon-stimulated gene involved in the control of protein translation and inflammation. J Biol Chem 289, 26642–26657.

Witkos, T.M., and Lowe, M. (2015). The Golgin Family of Coiled-Coil Tethering Proteins. Front Cell Dev Biol 3, 86.

Wong, M., Gillingham, A.K., and Munro, S. (2017). The golgin coiled-coil proteins capture different types of transport carriers via distinct N-terminal motifs. BMC Biol 15, 3.

Wong, M., and Munro, S. (2014). Membrane trafficking. The specificity of vesicle traffic to the Golgi is encoded in the golgin coiled-coil proteins. Science 346, 1256898.

Yeh, T.Y., Sbodio, J.I., Tsun, Z.Y., Luo, B., and Chi, N.W. (2007). Insulin-stimulated exocytosis of GLUT4 is enhanced by IRAP and its partner tankyrase. Biochem J 402, 279–290.

